# Ovine CD14- an immune response gene has a role against gastrointesinal nematode *Haemonchus contortus* -a novel report

**DOI:** 10.1101/2021.02.04.429682

**Authors:** Kavita Rawat, Aruna Pal, Samiddha Banerjee, Abantika Pal, Subhas Chandra Mandal, Subhashis Batabyal

**Author notes:** Both the authors contributed equally (Kavita Rawat, Aruna Pal). **Corresponding Author** & Assistant Professor (Animal Genetics and Breeding), West Bengal University of Animal and Fishery Sciences. 37, K.B.Sarani, Kolkata-37 (Former Scientist C, Department of Biotechnology, Ministry of Science and Technology, Govt. of India). Corresponding Author: Dr. Aruna Pal, M.V.Sc, PhD. Assistant Professor (Animal Genetics and Breeding), West Bengal University of Animal and Fishery Sciences, 37, KB Sarani, Kolkata-37, West Bengal. Pin-700037, E mail. Ph no.91- 9432349154. KR has conducted the experimental work, collected samples, performed acquisition of data, drafted the article. *ArP has conceptualized and designed the work, conducted research, analysed and interpreted the data, and drafted the article. SaB has conducted the experimental work. AbP has conducted the bioinformatics analysis and its interpretations.SCM has guided through the experiments on parasitology. SuB has conceptualized the work, guided through the biochemical analysis.

## Abstract

CD14 (also known as monocyte differentiation antigen) is an important immune response gene known to be primarily responsible for innate immunity against bacterial pathogen and as pattern recognition receptor (PRR) binds with LPS (endotoxin), lipoproteins, lipotechoic acid of bacteria.So far very limited work has been conducted in parasitic immunology. In the current study, we reported the role of CD14 in parasitic immunology in livestock species (sheep) for the first time. Ovine CD14 is characterized as a horse-shoe shaped as a bent solenoid with a hydrophobic amino-terminal pocket for CD14 along with domains. High mutation frequency was observed, out of total 41 mutations identified, 23 mutations were observed to be thermodynamically unstable and 11 mutations were deleterious in nature, causing major functional alteration of important domains of CD14, an indicative for variations in individual susceptibility for sheeps against *Haemonchus contortus* infestations. *In silico* studies with molecular docking reveals a role of immune response against *Haemonchus contortus* in sheep, which is later confirmed with experimental evidence through differential mRNA expression analysis for sheep, which revealed better expression of CD14 in *Haemonchus contortus* infected sheep compared to that of healthy sheep. We confirmed the above findings with supportive evidences with haematological and biochemical analyses. Phylogenetic analysis was conducted to assess the evolutionary relationship with respect to human and observed that sheep may well be used as model organism due better genetic closeness compared to that of mouse.

## 1. Introduction

CD14 molecule is an important immune response molecule and acts as pattern recognition receptor for innate immunity reported mostly against bacteria. In human, there are various types of CD molecules ranging from 1 to 371 (Engel et al., 2015). CD14 exists as both membranous (mCD14) and soluble (sCD14) form (Bazil et al., 1986). mCD14 is GPI-anchored and is mostly expressed on neutrophils, macrophages, and monocytes, and less expressed on a variety of other stromal and hematopoietic cells (Hazoit et al., 1988, Simmons et al., 1989). Three basic mechanisms are responsible for formation of sCD14, these are bypassing of GPI addition, cleavage of the GPI anchor by phospholipase D, or direct proteolytic cleavage from the cell surface (Wright et al, 1990). Expression of soluble CD14 is observed in serum, cerebrospinal, and other body fluids. CD14 is a pattern recognition receptor (PPR) responsible for the innate immune response to infection through the process of sensitization of host cells to bacteria. Bacterial lipopolysaccharide or LPS (endotoxin), lipoproteins, lipoteichoic acid and other acylated microbial products are the site of action for CD14. CD14 acts through various TLR (Toll-like Receptor) signalling complexes which in turn upon ligand binding induce intracellular proinflammatory signalling cascades (Goldsby et al., 2000).

In Gram-negative bacteria, the outer membrane consist of mostly lipopolysaccharide (LPS) (Rietschel et al., 1985). LPS-binding protein (LBP) aids in the process of binding of LPS to CD14 (Hailman et al., 1994, Tobias et al., 1995), is a 60-kDa serum glycoprotein produced by liver hepatocytes (Tobias et al., 1986, Schumann et al., 1990, Yu and Wright, 1996). With the aid of LBP, CD14 binds monomeric LPS (Hailman et al., 1994) and combines to the TLR4 complex. This in turn ultimately concentrates low levels of LPS, leading to the increase of the sensitivity of the system (Hailman et al., 1994, Tobias et al., 1995, Goyert et al., 1988, Pugin et al., 1994). MD-2 is another important immune molecule of molecular weight of 25-kD, which acts as a co-receptor is physically associated with TLR4 and is the major LPS acceptor of the TLR4 receptor complex. The binding of LPS to the TLR4–MD-2 complex induces a homodimeric receptor signaling complex comprises of two units each of TLR4 monomers, MD-2 proteins, and LPS molecules (Shimazu et al., 1999). Intracellular signaling domain of TLR4 is responsible for cell signaling, which leads to intracellular signaling adaptors after dimerization (Nunez Miguel et al., 2007). Small gel filtration mixing studies have revealed the role of CD14 as a catalyst in the process of transportation of LPS to the TLR4–MD-2 complex (Goyert et al., 1988, Gioannini, 2004, Prohinar et al., 2007, Teghanemt et al., 2007, Teghanemt et al., 2008).

Other than LPS, the CD14 molecule has the potential to bind a wide range of natural and synthetic acylated ligands due to its pattern recognition characteristics. For instance, CD14 and LBP combine together and causing opsonisation of whole bacteria and apoptotic cells leading to the clearance of infection and reduction of inflammation (Jack et al.,1995, Grunwald et al., 1996, Fan et al., 1999, Stelter et al., 1997, Schiff et al., 1997). CD14 is also capable to interact and bind to certain host phospholipids (Wurfel and Wright, 1997, Yu et al., 1997, Wang et al., 1998). Further, CD14 had been shown to bind to a wide range of acylated bacterial agonists of TLR2, namely lipoteichoic acid, peptidoglycan, mycobacterial lipoarabinomannan, atypical LPSs, and lipoproteins increasing cellular inflammatory responses (Nakata et al., 2006, Dziarski et al., 1998, Gupta et al., 1996, Savedra et al., 1996, Yu et al., 1998, Elass et al., 2007, . Lien et al., 1999, Takeuchi et al., 1999, Salomao et al, 2012, Opal and Cohen, 1999, Kim et al., 2005). It has been observed that soluble CD14 induces B cell growth and differentiation as present in bovine colostrums and milk (Fillip et al., 2001).

Thus it is evident that CD14 is an immune response molecule against both gram negative and gram positive bacteria (such as Mycobacterium sp., Pseudomonas sp. and Staphylococcus aureus). CD14 is an important drug target for the treatment of various diseases, like sepsis (Salomao et al., 2012, Opal and Cohen 1999), mastitis (Lee et al., 2003), treponemiasis (Schroder et al., 2000) and glomerulonephritis (Yoon et al., 2003), since it has a role in TLR agonist delivery. CD14 is equally important for parasitic diseases (Passos et al., 2017, Parija, S.C. 2011, Maizels, R.M. 2009, Das and Laha, 2017). CD14 as immune response gene may be utilized as various modes for disease management. These include therapeutic use of recombinant protein, gene therapy with cloned insert, evolvement of transgenic or gene edited disease resistant animals.

Genes for innate resistance may be of practical importance for studying phylogenetic relationship. Phylogenetic analysis has been carried out for different species including mammals and avian from time to time to find out the evolutionary relationship at the nucleotide level. Sequencing of genomes provides better opportunities for enriching our basic knowledge to the evolutionary pathways of related species. Mutation and natural selection are the driving force for the current day genome of each species arising through millions of years. Some preliminary comparisons between humans and other mammals have already been made (Hixson and Brown, 1986, Vigilant et al., 1991). Since CD14 gene shows a considerable degree of variability among the species, it can be used as a marker for phylogenetic analysis.

Cloning and sequencing of CD14 gene in domestic ruminants like cattle (*Bos taurus*, Pal et al.,2011), goat (*Capra hircus*, Pal et al., 2009, Pal et al., 2014), Buffalo (*Bubalus bubalis*, Pal et al., 2014), have already been conducted earlier in our lab. The present investigation aims for cloning and sequencing of the CD14 gene in sheep (*Ovis aries*), another important domestic ruminant. Currently, only the crystal structure of mouse CD14 (unliganded) purified from SF9 insect cells (Kim et al., 2005) and human CD14 (Kelley et al., 2013) are available in data base. Both mouse and human CD14 have an N-terminal hydrophobic cavity that provides a putative binding site for LPS and other acylated ligands. So, the present work objective is to computationally model the structures of CD14 in sheep using the existing experimentally determined structures of CD14 of livestock we studied earlier. Additionally, it focuses to identify and compare structural and functional domain of CD14 derived peptide using bioinformatics tools and to explore the domains in 3D structure responsible for pathogen binding. The molecular phylogenetic analysis in domestic ruminants with other domestic and wild mammal and avian with respect to CD14 gene (responsible for innate resistance) and derived peptide, has also been studied which might be helpful to look into in evolutionary ecology and pharmacogenomic studies. *In silico* analysis with molecular docking were conducted to detect the role of CD14 in parasitic immunity. Differential mRNA expression profiling for healthy and diseased (Haemonchus infested) sheep with respect to CD14 gene was conducted to confirm above findings.

## 2. Materials and methods

### Sample Collection and RNA Isolation

Sheep abomassal tissue (1g) was collected from slaughter house located under Kolkata Municipality Corporation. Adult males (n=6) of about 1-1.5 years of age were selected for collection of samples. Tissue was immersed in Trizol in vial and transported in ice to the laboratory for RNA isolation. Total RNA was isolated using TRIzol extraction method (Life Technologies, USA), as per the standard procedure and was further utilized for cDNA synthesis (Pal and Chatterjee, 2009; Pal et al., 2011). Since the samples were collected from the slaughter house, ethical approval was not necessary. cDNA concentration was estimated and samples above 1200 microgram per ml were considered for further study.Tissue from abomassum was collected from both healthy and sheep infected with *Haemonchus contortus* for studying the expression profiling with quantitative PCR. Abomassal tissue was collected from both tip and middle of the abomassum. Expression from tip of abomassum was considered as control during ddct calculation.

### cDNA synthesis and PCR Amplification of the CD14 Gene

20μL of reaction mixture was composed of 5μg of total RNA, 0.5μg of oligo dT primer (16–18mer), 40U of Ribonuclease inhibitor, 1000M of dNTP mix, 10mM of DTT, and 5U of MuMLV reverse transcriptase in appropriate buffer. The reaction mixture was mixed thoroughly followed by incubation at 37°C for 1 hour. The reaction was allowed upto 10 minutes by heating the mixture at 70°C unliganded and then chilled on ice. Afterwards, the integrity of the cDNA was checked by performing PCR. Concentration of cDNA was estimated through Nanodrop. Ovine CD14 primer pair was designed based on the CD14 mRNA sequences of cattle (GenBank Acc No.AF141313) using DNASTAR software (Hitachi Miraibio Inc., USA) to amplify full length open reading frame (ORF) of CD14 gene sequence of sheep. The primers were Forward: CD14-1-F ATGGTCTGCGTGCCCTACCTG and Reverse: CD14-70-1- R GGAGCCCGAGGCTTCGCGTAA. 25μL of reaction mixture was comprised of 80–100ng cDNA, 3.0μL 10X PCR assay buffer, 0.5μL of 10mM dNTP, 1U Taq DNA polymerase, 60ng of each primer, and 2mM MgCl_2_. PCR-reactions were performed in a thermocycler (PTC-200, MJ Research, USA) with cycling conditions as, initial denaturation at 94°C for 3min, further denaturation at 94°C for 30sec, annealing at 61°C for 35sec, and extension at 72°C for 3min were conducted for 35 cycles followed by final extension at 72°C for 10 min.

### cDNA Cloning and Sequencing

Amplified product of ovine CD14 was checked by 1% agarose gel electrophoresis. The products were purified from gel using Gel extraction kit (Qiagen GmbH, Hilden, Germany) .pGEM-T easy cloning vector (Promega, Madison, WI, USA) was used for cloning. Then, 10μL of ligated product was mixed thoroughly to 200μL competent cells, and heat shock was given at 42°C for 45sec in a water bath. Subsequently, the cells were immediately transferred on chilled ice for 5 min., and SOC medium was added to it. The bacterial culture was centrifuged to obtain the pellet and plated on LB agar plate containing Ampicillin (100mg/mL) added to agar plate @1:1000, IPTG (200mg/mL) and X-Gal (20mg/mL) for blue-white screening. Plasmid isolation from overnight-grown culture was carried out by small-scale alkaline lysis method as described in (Sambrook et al., 2001). Recombinant plasmids were characterized by PCR using CD14 primers as reported earlier and restriction enzyme digestion. CD14 gene fragments released by enzyme EcoRI (MBI Fermentas, USA) were inserted in recombinant plasmid which was sequenced by dideoxy chain termination method with T7 and SP6 primers in an automated sequencer (ABI prism, Chromous Biotech, Bangalore).

### Sequence Analysis

DNASTAR Version 4.0, Inc., USA software was employed for the nucleotide sequence analysis for protein translation, sequence alignments, and contigs comparisons. Novel sequence was submitted to the NCBI Genbank and accession number KY110723.1 was obtained which is available in public domain. CD14 sequences have already been analyzed earlier in our lab for cattle (GU368102), buffalo (DQ457089) and goat (DQ457090).

### Study of Predicted ruminant CD14 Protein Using Bioinformatics Tools

Predicted peptide sequences of CD14 gene of sheep and other ruminants (cattle, buffalo, goat), characterized in our lab earlier, were then aligned with the CD14 peptide of other rodent species and *Homo sapiens* using MAFFT (Katoh and Standley, 2016). The analysis was conducted for a sequence based comparative study among ruminants with rodent and human. Signal peptide is essential to prompt a cell to translocat<u>e</u> the protein, usually to the cellular membrane and ultimately signal peptide is cleaved to give mature protein. Prediction of presence and location of signal peptide of CD14 gene was conducted using the software (SignalP 3.0 Sewer-prediction results, Technical University of Denmark). Pattern recognition receptor as CD14 is rich in Leucine, hence it is essential to calculate leucine percentage. Leucine percentage was calculated manually from predicted peptide sequence. Di-sulphide bonds are essential for protein folding and stability, ultimately. It is the 3D structure of protein which is biological active. Di-sulphide bonds were predicted using suitable software (http://bioinformatics.bc.edu/clotelab/DiANNA/) (Kim et al., 2005).

Protein sequence level analysis was employed (http://www.expasy.org./tools/blast/) for assessment of leucine rich repeats (LRR), leucine zipper, detection of Leucine-rich nuclear export signals (NES), and detection of the position of GPI anchor, N-linked glycosylation sites. Since CD14 is a pattern recognition receptor, these are rich in leucine rich repeat, which are essential for pathogen recognition and binding. Leucine zipper is essential for dimerization of CD14 molecule. N-linked glycosylation is important for the molecule to determine its membranous or soluble form. Leucine rich nuclear export signal is essential for export of CD14 from nucleus to cytoplasm, whereas GPI anchor is responsible for anchoring in case of membranous protein. Prosite was used for LRR site detection.

Leucine-rich nuclear export signals (NES) was analyzed with NetNES 1.1 Server, Technical university of Denmark. O-linked glycosylation sites were detected using NetOGlyc 3.1 server (http://www.expassy.org/), whereas N-linked glycosylation site were assessed through NetNGlyc 1.0 software (http://www.expassy.org/). Sites for leucine-zipper were detected through Expassy software, Technical university of Denmark (Glick, 1977). Sites for alpha helix and beta sheet were detected using NetSurfP-Protein Surface Accessibility and Secondary Structure Predictions, Technical University of Denmark (Petersen et al., 2009). Domain linker site were predicted (Ebina et al., 2009). LPS-binding (Cunningham et al., 2000) and LPS-signalling sites (Muroi et al., 2002) were predicted based on homology studies with other species CD14 polypeptide. These sites are important for pathogen recognition and binding.

### Three dimensional structure prediction and Model quality assessment

Three dimensional model of ovine CD14 polypeptide was predicted through Swissmodel repository (Kiefer et al., 2009). Templates possessing greatest identity of sequences with our target template were identified with PSI-BLAST (http://blast.ncbi.nlm.nih.gov/Blast, Altschul et al., 1997). PHYRE2 server based on ‘Homology modelling approach’ was used to build three dimensional model of ovine CD14 (Kelley, 2015). Molecular visualization tool as PyMOL (http://www.pymol.org/) was employed for model generation and visualization of three dimensional structure of ovine CD14. The structure of ovine CD14 molecule was evaluated and assessed for its stereochemical quality (through SAVES, Structural Analysis and Verification Server, http://nihserver.mbi.ucla.edu/SAVES/); then refined and validated through ProSA (Protein Structure Analysis) web server (https://prosa.services.came.sbg.ac.at/prosa) (Wiederstein and Sippl, 2007). NetSurfP server (http://www.cbs.dtu.dk/services/NetSurfP/ Peterson et al., 2009) was used for assessing surface area of ovine CD14 through relative surface accessibility, Z-fit score, and probability of alpha-Helix, beta-strand and coil score.

Alignment of 3-D structure of ovine CD14 with other ruminant species as cattle, buffalo and goat were analyzed with RMSD estimation to evaluate the structural differentiation by TM-align software (Zhang et al., 2005). Thermodynamic stability of the protein with mutations and the deleterious nature of the mutant amino acid were analyzed through I-mutant 2.0 (Capriotti et al., 2005) and Provean analysis (Choi and Chan, 2015) respectively.

### Protein-protein interaction network depiction

In order to understand protein interaction network of CD14 protein, we performed search in STRING 9.1 database (Franceschini et al., 2015). The functional interaction was assessed with confidence score. Interactions with score < 0.3, scores ranging from 0.3 to 0.7, and scores >0.7 are classified as low, medium and high confidence respectively. Also, we executed KEGG analysis which depicted the functional association of CD14 with other related proteins.

### Phylogenetic analysis

Nucleotide sequence of Ovine CD14 was aligned with the published nucleotide sequences of other ruminant species we characterized earlier and also with other species obtained from gene bank (http://www.ncbi.nlm.nih.gov/blast) as listed in **Table 1** for CD14. ClusterIV method of MegAlign Programme of Lasergene Software (DNASTAR) was employed for the analysis.

**Table 1:**
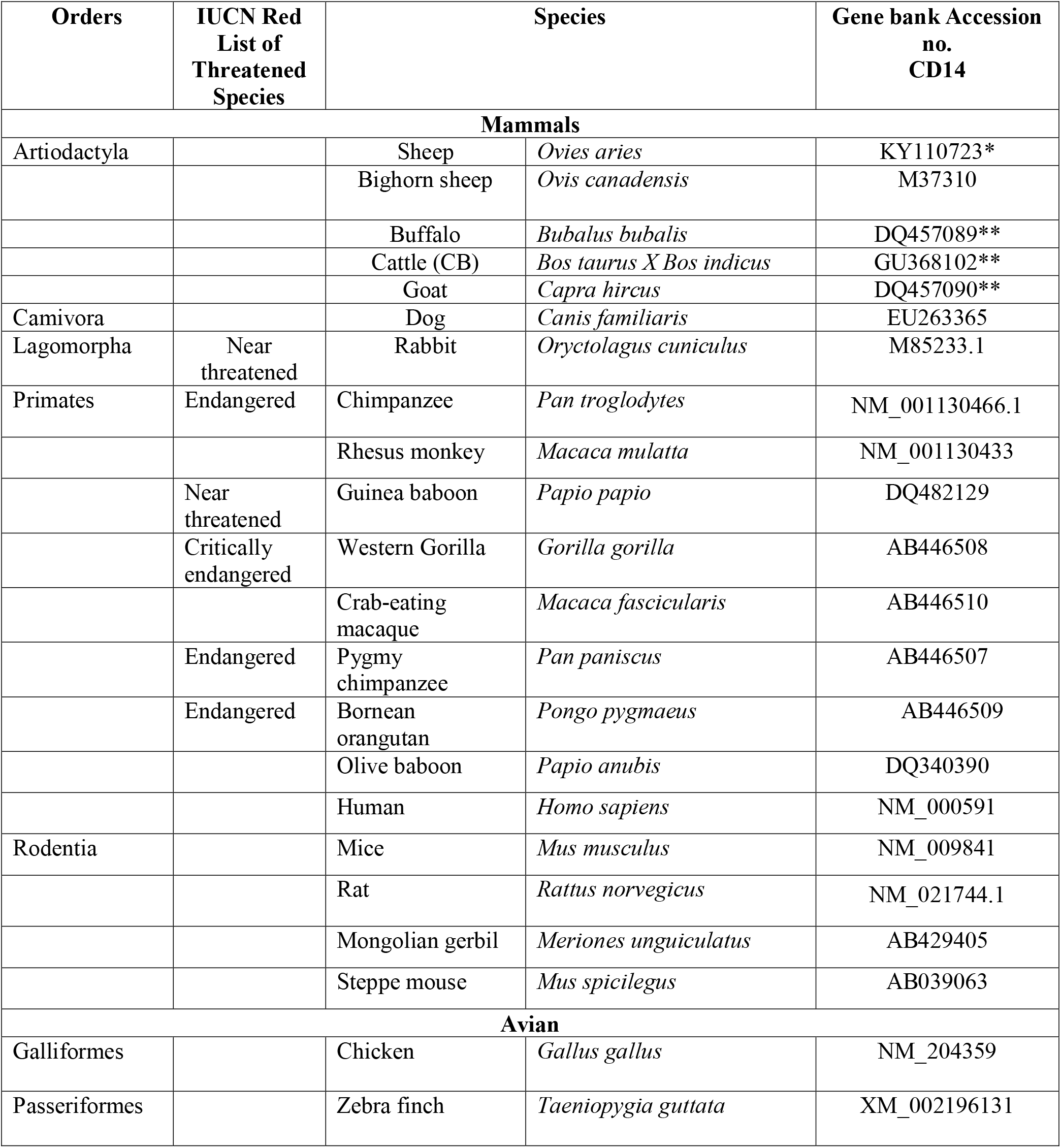
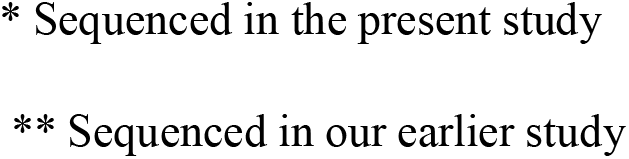
List of mammals & avian under present study.

| Orders | IUCN Red List of Threatened Species | Species |  | Gene bank Accession no. CD14 |
| --- | --- | --- | --- | --- |
| Mammals |  |  |  |  |
| Artiodactyla |  | Sheep | *Ovies aries* | KY110723* |
|  |  | Bighorn sheep | *Ovis canadensis* | M37310 |
|  |  | Buffalo | *Bubalus bubalis* | DQ457089** |
|  |  | Cattle (CB) | *Bos taurus X Bos indicus* | GU368102** |
|  |  | Goat | *Capra hircus* | DQ457090** |
| Camivora |  | Dog | *Canis familiaris* | EU263365 |
| Lagomorpha | Near threatened | Rabbit | *Oryctolagus cuniculus* | M85233.1 |
| Primates | Endangered | Chimpanzee | *Pan troglodytes* | NM_001130466.1 |
|  |  | Rhesus monkey | *Macaca mulatta* | NM_001130433 |
|  | Near threatened | Guinea baboon | *Papio papio* | DQ482129 |
|  | Critically endangered | Western Gorilla | *Gorilla gorilla* | AB446508 |
|  |  | Crab-eating macaque | *Macaca fascicularis* | AB446510 |
|  | Endangered | Pygmy chimpanzee | *Pan paniscus* | AB446507 |
|  | Endangered | Bornean orangutan | *Pongo pygmaeus* | AB446509 |
|  |  | Olive baboon | *Papio anubis* | DQ340390 |
|  |  | Human | *Homo sapiens* | NM_000591 |
| Rodentia |  | Mice | *Mus musculus* | NM_009841 |
|  |  | Rat | *Rattus norvegicus* | NM_021744.1 |
|  |  | Mongolian gerbil | *Meriones unguiculatus* | AB429405 |
|  |  | Steppe mouse | *Mus spicilegus* | AB039063 |
| Avian |  |  |  |  |
| Galliformes |  | Chicken | *Gallus gallus* | NM_204359 |
| Passeriformes |  | Zebra finch | *Taeniopygia guttata* | XM_002196131 |

\* Sequenced in the present study
\*\* Sequenced in our earlier study

### Differential mRNA expression profiling for CD14 gene with respect to healthy and diseased sheep

Sheep population under current study was divided in two groups, as diseased (infected with parasitic infection of *Haemonchus contortus*) and healthy (no infection), as assessed through egg per gram count.

### Animals, faecal sample collection and determination of FEC

We had randomly collected 60 sheep samples from LFC farm,WBUAFS, Mohanpur campus. The samples were collected prior to routine deworming procedure, presumed to have pre existing gastro intestinal parasite since they were not dewormed for last three months. The animals were presumed to get exposed to natural infection during grazing.

Faecal samples of the sheep were examined thoroughly by salt floatation method and the Faecal Egg Count were screened for each and every sample. Based on statistical analysis, samples were collected from two groups, designated as healthy (x◻ +SD, where x◻ stands for mean and SD means standard deviation) and diseased (x◻− SD, where x◻ stands for mean and SD means standard deviation). Sheep were regularly sold and slaughtered for the purpose of mutton production unit. Altogether 12 tissue samples were screened, 6 each with high FEC (grouped as diseased) and 6 each with low FEC (grouped as healthy) for further case study. Tissue samples were collected for abomassum, rumen, small intestine, caecum, liver and lymph node.

Serum was harvested from blood samples as per the standard procedure. The serum biochemical parameters like glucose, total protein, albumin, globulin, uric acid, creatinine, BUN, triglyceride, cholesterol, HDL, LDL concentrations and serum enzyme variables (AST, ALT, & AP) were estimated by using a semi-auto biochemistry analyzer (Span diagnostic Ltd.) with standard kits.

### Collection of blood samples

Blood sample was collected aseptically from the juglar vein of sheep in the separate sterile vials with anticoagulants in the morning hours between 8 a.m. to 10 a.m. 2-3ml blood was used for hematological parameters, 5ml blood for serum without anticoagulant were collected centrifuged (10 min at 1000 rpm) and preserved at − 20 °C until analysis for other biochemical and serological tests.

#### Haematological Profiles

The haematological parameters like hemoglobin, erythrocyte sedimentation rate (ESR) and packed cell volume (PCV) were estimated in whole blood soon after the collection of blood. Hemoglobin was estimated by acid haematin method (Benjamin, 1985), E.S.R. and PCV by Wintrobe’s tube (Hawk, 1965). The total erythrocyte count (TEC), total leukocyte count (TLC) and Differential leukocyte count (DLC) were studied by standard methods described by Jain (1986).

#### Biochemical Analysis

The serum biochemical parameters, estimated in the experiments were total protein, albumin, globulin, albumin: globulin, aspartate aminotransferase (AST), alanine aminotransferase (ALT), alkaline phosphatase (ALP), Total bilirubin, Indirect bilirubin, direct bilirubin, glucose, uric acid, urea and BUN by using a semi-auto biochemistry analyzer (Span diagnostic Ltd.) with standard kits (Trans Asia Bio-Medicals Ltd., Solan, HP, India). The methodology used for estimation of total protein, albumin, total & direct bilirubin, ALT, ALP, glucose, creatinine urea and uric acid were biuret method, bromocresol green (BCG) method, 2-4-DNPH method, modified kind and king’s method, GOD/POD method, modified Jaffe’s Kinetic method. GLDH-urease method and trinder peroxidise method respectively.

### Real Time PCR (qRT-PCR)

Equal quantity of cDNA as quantified through Nanodrop was used in each reaction of 96 well optical plate of ABI 7500 system. Each reaction consisted of 1ng cDNA template, 10 μl of 2X SYBR Green PCR Master Mix, 20pMol each of forward and reverse primers and make up final volume upto 20 μl with nuclease free water. Each sample was run in triplicate. Analysis of real-time PCR (qRT-PCR) was performed by delta-delta-Ct (ΔΔCt) method, Ct denotes the threshold value. The list of primers used for QPCR study have been listed below with annealing temperature at 60 C.

CD14 primer:

F ACCACCCTCAGTCTCCGTAA,
R-GTGCTTGGGCAATGTTCAG

18S rRNA primer:

F-TCCAGCCTTCCTTCCTGGGCAT,
R-GGACAGCACCGTGTTGGCGTAGA.

#### *In silico* study for the detection of binding site of CD14 with *Haemonchus contortus* with Molecular Docking

Molecular docking is a bioinformatics tool used for *in silico* analysis for the prediction of binding mode of a ligand with a protein 3D structure. Patch dock is an algorithm for molecular docking based on shape complementarity principle (Duhovny et al., 2005). Patch Dock algorithm was used to predict ligand protein docking for surface antigen for gastrointestinal parasitism with *Haemonchus contortus* (alpha tubulin and beta tubulin antigen) with ovine CD14.

## 3. Result

### Cloning and sequencing of CD14 cDNA of sheep

The ovine CD14 was observed to have an open reading frame of 1116 nucleotide, ATG as start codon and TAA as stop codon (KY110723) (Fig 1). Nucleotide sequences as well as derived peptide sequences for *Bos taurus* (GU368102), *Bubalus bubalis* (DQ457089), and *Capra hircus* (DQ457090) had been sequenced earlier in our lab and submitted to the gene bank. Ovine CD14 derived peptide was observed to have 373 amino acids precursor corresponding to the coding sequence of the gene and a signal peptide consisting of 20 amino acids (Fig 2). The molecular weight of sheep CD14 was observed to be 39645 Dalton. Slight differences have been observed in the molecular weight of the CD14 molecule of ruminants.

**Fig 1:**
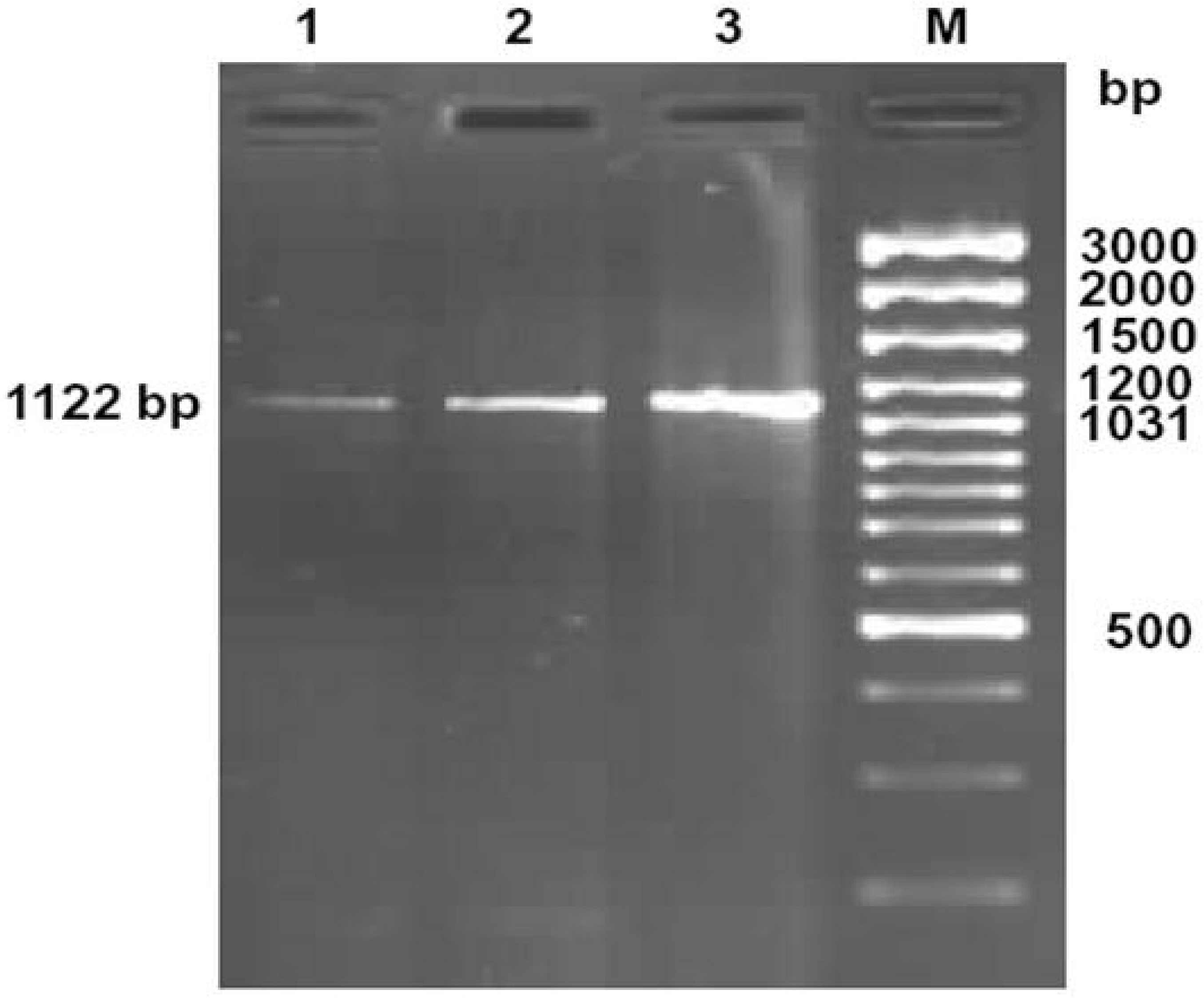
Amplification of 1122bp fragment of CD14 gene of sheep and goat. Lane 1-2: Amplified product from cDNA of CD14 gene of sheep Lane M: 100bp DNA ladder plus

**Fig 2:**
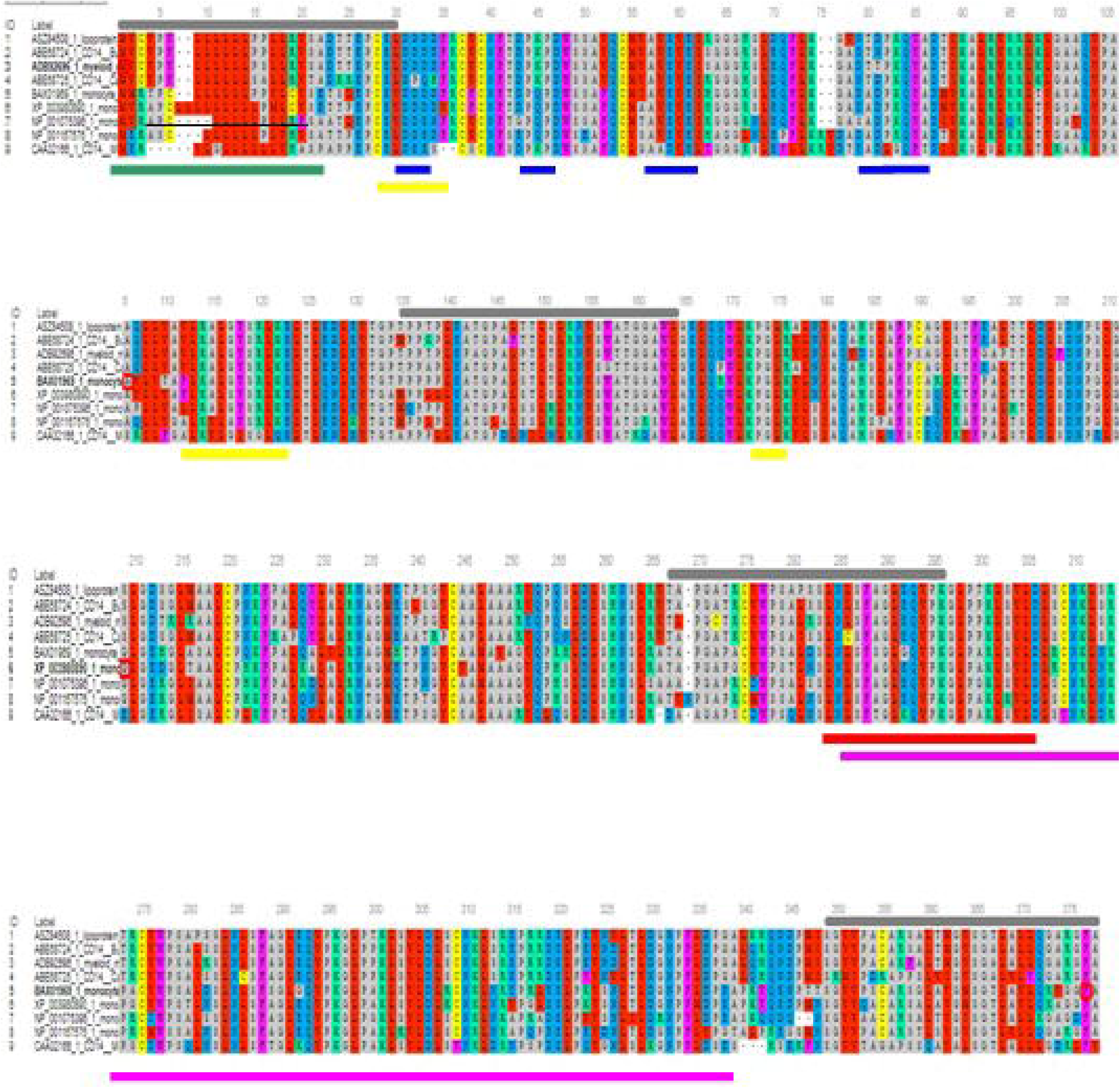
Alignment of ruminant CD14 derived peptide with other species. 1: Sheep (Ovis aries), 2: CB cattle (Bos taurus x Bos indicus), 3:Buffalo ( Bubalus bubalis), 4: Goat (Capra hircus), 5: Human (Homo sapiens), 6: Mice (Mus musculus), 7: Dog (Canis lupus familiaris), 8: Cat (Felis catus), 9: Horse (Equus caballus) Green line: Signal peptide Blue line: LPS binding site Yellow line: LPS signaling site Red line: Leucine zipper Black overline: Leucine rich repeat Red line: Leucine zipper Pink line: Domain linker prediction site

### Bioinformatics analysis for CD14 gene

Ovine CD14 was observed to have high GC content. GC content of CD14 gene in ruminants were found to be high (62.2 to 62.7%) depicted in Table 2. Alignment of the derived peptide sequence of ruminants have been depicted in Fig 4. Amino acid cysteine contains sulphur, which is responsible for disulphide bond formation, which in turn are responsible for protein folding. The site for disulphide bond has been predicted in Table 2. LPS binding and LPS signaling sites are very important to determine the potentiality for innate immunity of CD14 gene. Derived peptide of the ruminant CD14 gene differs in putative N-linked glycosylation sites at amino acid positions, 4 sites in cattle and buffalo and 3 sites in goat were predicted (Table2). O-linked glycosylation site for ruminant CD14 differs, three in Buffalo and five in both goat and crossbred cattle (Table 2), but absent in human [Savedra et al., 1996]. O-linked Glycosylation is needed when hydrophilic clusters of carbohydrates alter the polarity and solubility of protein or protein folding. CD14 and other Toll-like receptors were characterized by presence of LRR and high leucine content. Leucine rich repeats (LRR) predicted peptide sequence of ruminant CD14 cDNA were studied (Table 2, Fig 2). Protein dynamics study revealed that in ruminants, position of glycosyl phosphatidyl inositol (GPI) anchor located at C-terminus of the CD14 molecule differs (Table 2). Seven sites for leucine rich nuclear export signal (NES) were detected in CD14 gene of ruminants (Table 2, Fig 2). In terms of secondary structure prediction, 7 regions of alpha helix and eleven regions of Beta strands with different amino acid position in ruminant CD14 molecule were predicted. (Table 2). Leucine zipper pattern was found at amino acid position 279^th^ in ruminant CD14 gene (Fig 2). The site for Domain linker for ovine CD14 site was predicted, which is almost similar to amino acid positions in other ruminants (Table 2, Fig 2). 3-D model of CD14 molecule of sheep was predicted to be horseshoe shaped structure, with alternating alpha helix and beta chains in swiss model (Fig 3A), pymol view with LPS binding site (Fig 3B), surface view with LPS binding site (Fig 3C). Fig 3D depicts the groove or pathogen binding site of ovine CD14 surface view. Predicted protein of CD14 molecule revealed a bent solenoid containing a hydrophobic amino terminal pocket. Alignment of 3D structure of ovine CD14 with goat (Fig 4A), cattle (Fig 4B) and buffalo (Fig 4C) have been depicted. The variability in the structure of pocket of CD14 and alternative rim residues believed to be important for LPS binding and cell activation, leading to variation in innate immunity across the species. Pocket or groove (LPS binding site) for TM aligned CD14 sheep and CD14 buffalo have been depicted in Figure 5A. Similarly Fig 5B depicts pathogen binding sites for ovine CD14 with that of goat CD14. The amino terminal pocket containing grooves are involved ligand binding (Fig 9c). CD14 receptor exists as dimer with the help of leucine zipper in its biologically active form. It had been observed that the amino terminal pocket for ruminant CD14 with grooves is the site for ligand binding, which is the most important factor for CD14. The rim residues of the CD14 hydrophobic pocket are the primary determinant for binding with LPS, or other acylated CD14 ligands. Positively charged residues as K71 and R72 are present on the rim and R80, K87, and R92 are present just outside the rim, where as hydrophobic residues as W45, F49, V52, F69, Y82, and L89 encircle the rim. These residues were observed to be the primary determinant for ligand binding.

**Table 2:** Bioinformatics analysis of CD14 derived peptide for domestic ruminants.

|  | *Bubalus bubalis* | cross of *Bos taurus* X *Bos indicus* | *Ovis aries* | *Capra hircus*, |
| --- | --- | --- | --- | --- |
| Gene Bank Protein id | ABE68724 | ADB92696.1 | <u>ASZ84508.1</u> | <u>ABE68725.1</u> |
| GC content of gene | 62.3 | 62.7 | 62.92 | 62.21%. |
| leucine rich repeats | Six | Nine | Three | Seven |
| leucine content, | 17.2% | 16.89% | 16.89% | 15.5% |
| putative N-linked glycosylation sites, | 4 | 4 | 4 | 3 |
| sites for O-linked glycosylation | **3 sites** | 5 sites for O- linked glycosylation at amino acid positions 128, 131, 134, 145, 147, | 6 sites<br>Thr 13,17,19,21<br>131, 237, | 5 O-linked glycosylation sites. amino acid positions 128, 131, 139, 144 and 145 |
| disulphide bridges | **Five** | Four<br>amino acid positions<br>26-37, 39-52, 186-216, 240-270 | Four | Four |
| LPS binding site | 4 sites,<br>29<sup>th</sup> -32<sup>nd</sup> , 44<sup>th</sup> -<br>47<sup>th</sup> , 55<sup>th</sup> -59<sup>th</sup><br>and 77<sup>th</sup> -83<sup>th</sup><br>amino acid<br>positions | four LPS binding sites<br>were depicted from<br>29<sup>th</sup> -32<sup>nd</sup> , 44<sup>th</sup> -47<sup>th</sup> ,<br>55<sup>th</sup> -59<sup>th</sup> and 77<sup>th</sup> -83<sup>th</sup><br>amino acid positions | four LPS<br>binding sites<br>were depicted<br>from 29<sup>th</sup> -32<sup>nd</sup> ,<br>44<sup>th</sup> -47<sup>th</sup> , 55<sup>th</sup> -<br>59<sup>th</sup> and 77<sup>th</sup> -<br>83<sup>th</sup> amino acid<br>positions | Four LPS binding<br>site from 29<sup>th</sup> -32<sup>nd</sup><br>, 44<sup>th</sup> -47<sup>th</sup> , 55<sup>th</sup> -<br>59<sup>th</sup> and 77<sup>th</sup> -83<sup>th</sup><br>amino acid<br>positions |
| LPS signaling sites | Three LPS | Three LPS signalling | Three LPS | three LPS |
|  | signalling sites were predicted for amino acid positions as 27<sup>th</sup>-33<sup>th</sup>, 109-119<sup>th</sup>, 169-171<sup>th</sup> | sites were predicted for amino acid positions as 27<sup>th</sup>-33<sup>th</sup>, 109-119<sup>th</sup>, 169-171<sup>th</sup> amino acid | signalling sites were predicted for amino acid positions as 27<sup>th</sup>-33<sup>th</sup>, 109-119<sup>th</sup>, 169-171<sup>th</sup> amino acid | signaling sites<br>Three LPS signalling sites were predicted for amino acid positions as 27<sup>th</sup>-33<sup>th</sup>, 109-119<sup>th</sup>, 169-171<sup>th</sup> |
| regions of alpha helix | **7 regions of alpha helix from amino acid** 4-12, 46-52, 65-75, 186-190, 241-244, 335-337 and 361-367and | five regions were detected for alpha helix conformation as 4-12, 67-73, 241-244, 343-346, 360-368 |  | five regions of alpha helix amino acid positions 4-18, 46-52, 81-84, 241-244 and 361-370 |
| regions of beta strand | 11 regions for beta strands were predicted from amino acid positions 119-121, 145-147, 173-175, 181-182, 197-199, 225-227, 252-254, 278-280, 299-301, 321-323 and 345-347 in bubaline CD14 molecule | eleven regions were identified for beta strand conformation as 60-61, 118-122, 145-147, 173-176, 181-182, 197-199, 225-227, 252-254, 278-280, 299-301, 321-324 amino acid positions |  | twelve regions of beta sheet were predicted amino acid positions 26-27, 34-37, 59-60, 118-122, 145-147, 173-176, 181-182, 197-199, 225-228, 252-254, 299-301, 321-323 |
| , leucine rich nuclear export signal | 7 detected at amino acid | 7 sites at amino acid positions 12, 15, 16, 17, 117, 122,127 | 7 sites at amino acid positions 12, 15, 16, 17, | seven amino acid positions. acid positions 15, 16, |
|  | positions 12, 15, 16, 17,117, 122, 127 |  | 117, 122,127 | 17, 20, 117, 122, and 127 of goat CD14 peptide |
| leucine zipper | amino acid position 279<sup>th</sup> to 300<sup>th</sup> in bubaline CD14 gene | amino acid position 279 of CB CD14 peptide | amino acid position 279 of CB CD14 peptide | Leucine zipper pattern was found at amino acid position 279 |
| domain linker, | Domain linker site was predicted for amino acid positions 121-146 and 283-334 | for amino acid positions 121-146 and 283-334 | for amino acid positions 121-146 and 283-334 | Domain linker site was predicted for amino acid positions 121-150 and 283-334. |
| site for a glycosyl phosphatidyl inositol anchor located at C-terminus | glycosyl phosphatidyl inositol (GPI) anchor located at C-terminus near 353<sup>th</sup> position of the CD14 molecule | glycosyl phosphatidyl inositol (GPI) anchor located at C-terminus near 345<sup>th</sup> position of the CD14 molecule | glycosyl phosphatidyl inositol (GPI) anchor located at C-terminus near 353<sup>th</sup> | glycosyl phosphatidyl inositol (GPI) anchor located at C-terminus near 344<sup>th</sup> position of the CD14 molecule |
| Molecular weight | 39705.07 Daltons | 39939.29Daltons | 39645.00 | 39929.42 Da |

**Fig 3:**
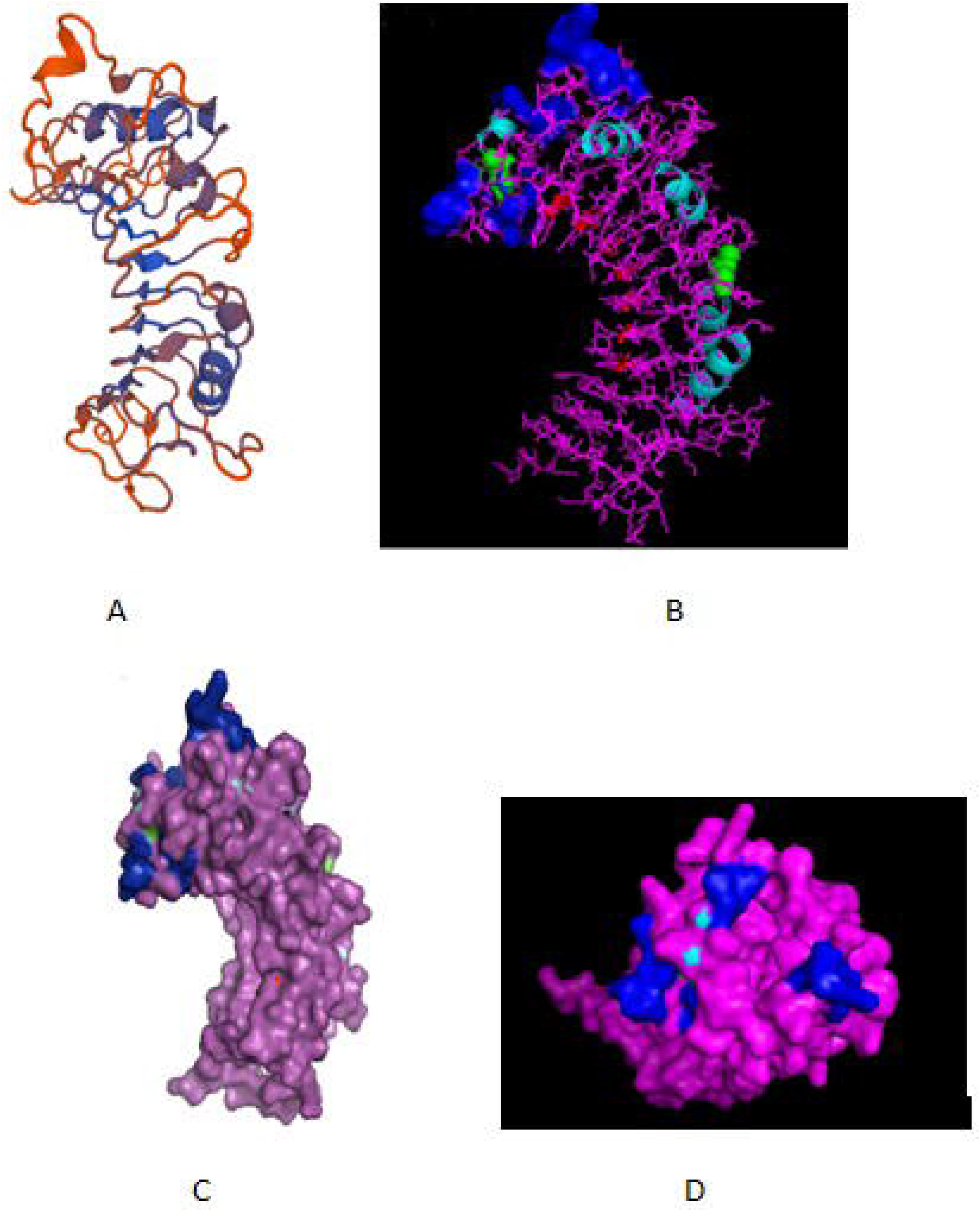
3D structure of sheep CD14. A: 3D structure of ovine CD14 gene-Swiss prot view **B:** 3D structure of ovine CD14 gene-Pymol view (with LPS binding sites) C: 3D structure of ovine CD14 gene-Surface view (with LPS binding sites) D: Sheep CD14- groove (pathogen binding site)

**Fig 4:**
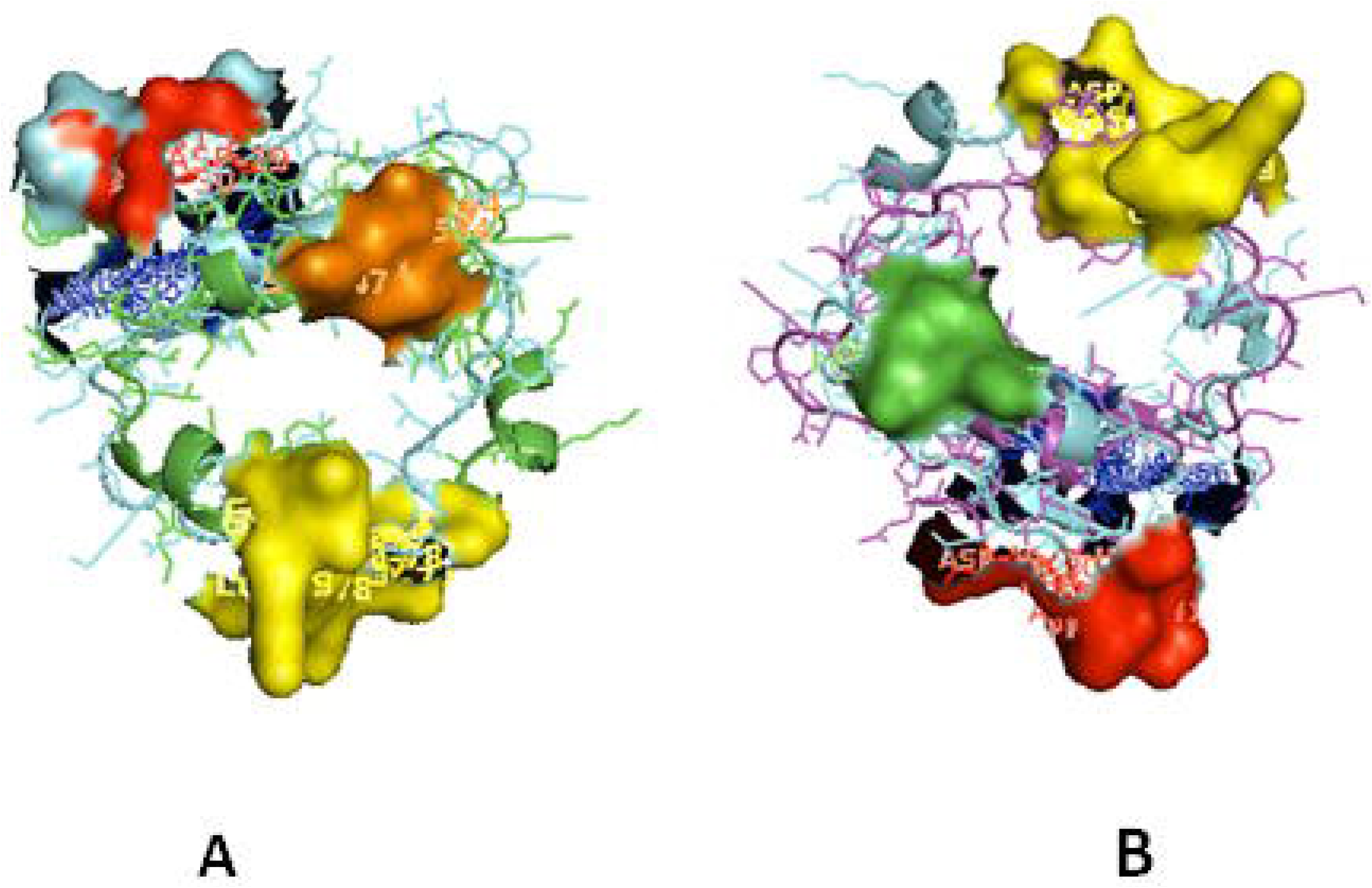
TM alignment of 3D structure of sheep with other ruminants. A: TM alignment of the 3D structure of sheep and goat CD14 molecules. Green: Sheep CD14 molecule, Cyan: Goat CD14 molecule B: TM alignment of the 3D structure of sheep and cattle CD14 molecules. Green: Sheep CD14 molecule, Orange: Cattle CD14 molecule C: TM alignment of the 3D structure of sheep and buffalo CD14 molecules. Green: Sheep CD14 molecule, Pink: Buffalo CD14 molecule

**Fig 5:**
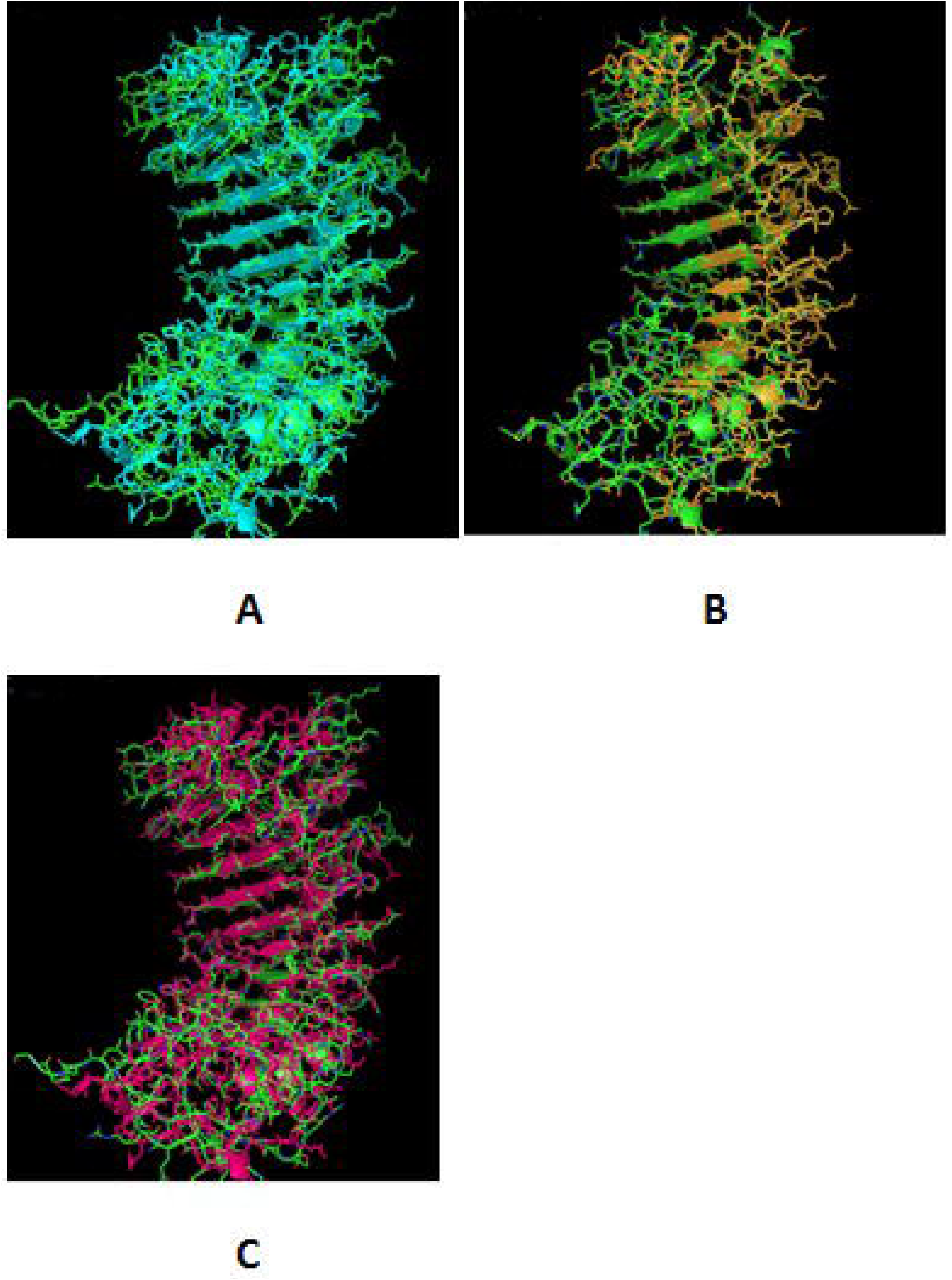
Depiction of receptor binding site for CD14. A: Pocket or groove (LPS binding site) for TM aligned CD14 sheep and CD14 buffalo B: Pocket or groove (LPS binding site) for TM aligned CD14 sheep and CD14 goat

### Comparison of ovine CD14 with goat

Small ruminants as sheep and goat were observed to have significant difference in susceptibility to commonly occurring diseases. Leucine content was observed to be more in sheep CD14 compared to that of goat. CD14 derived peptide of sheep were observed to have comparatively less GC content, but more leucine content percentage. Number of LRR in buffaloes were observed to be less in buffalo compared to cattle (Table 2). GPI anchor was present at 353^rd^ position in buffalo, in constrast to 345^th^ position of cattle. GPI anchor is important for membranous form of CD14 gene. 353^rd^ position implies greater availability of CD14.

Alignment of CD14 (3 D structure) of sheep/goat, sheep/ cattle, sheep/buffalo have been depicted in Fig 4A,Fig 4B and Fig 4C respectively. LPS binding sites were situated at the rim of the groove of the receptor. Four LPS binding sites have been depicted as 4 sites as 29^th^-32^nd^, 44^th^-47^th^, 55^th^-59^th^ and 75^th^-80^th^ amino acid positions and depicted in red, green, blue, yellow respectively (Fig 5A-B). Comparison of CD14 structure of Sheep/goat were observed to be highly superimposable with a root mean square deviation of 0.23 A °. Observation of goat/ buffalo CD14 were also observed to be greatly superimposable with RMSD being 1.89. The structural differences arising due to nucleotide differences have been depicted. Comparison of CD14 structure of Sheep/buffalo were observed to be highly super imposable with a root mean square deviation of 0.23. String analysis (Fig 6) and KEGG analysis (Fig7) depicted that CD14 works in combination with TLR4 and MD2.

**Fig 6:**
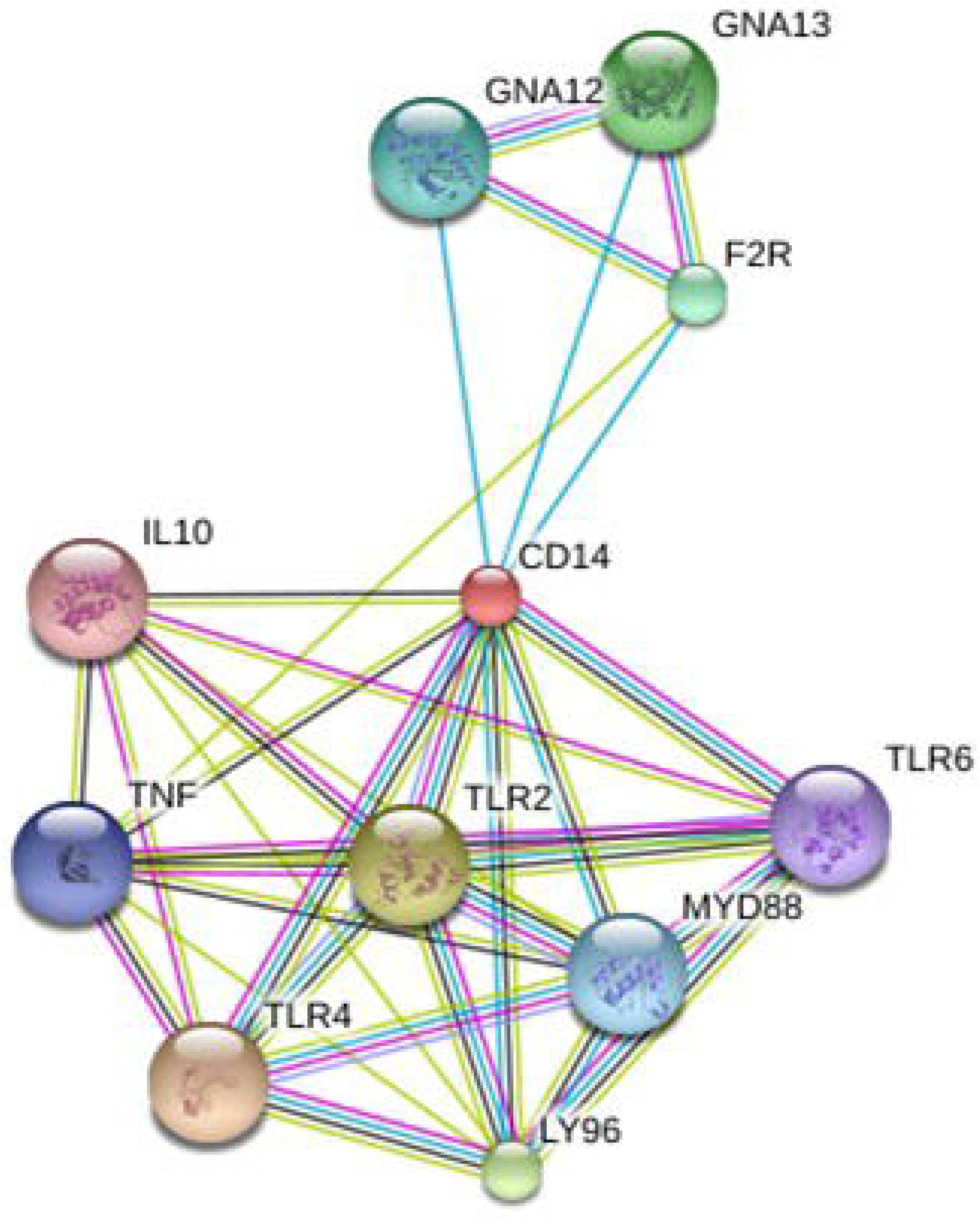
String network-Molecular interaction of ovine CD14 molecule with other related molecules

**Fig7:**
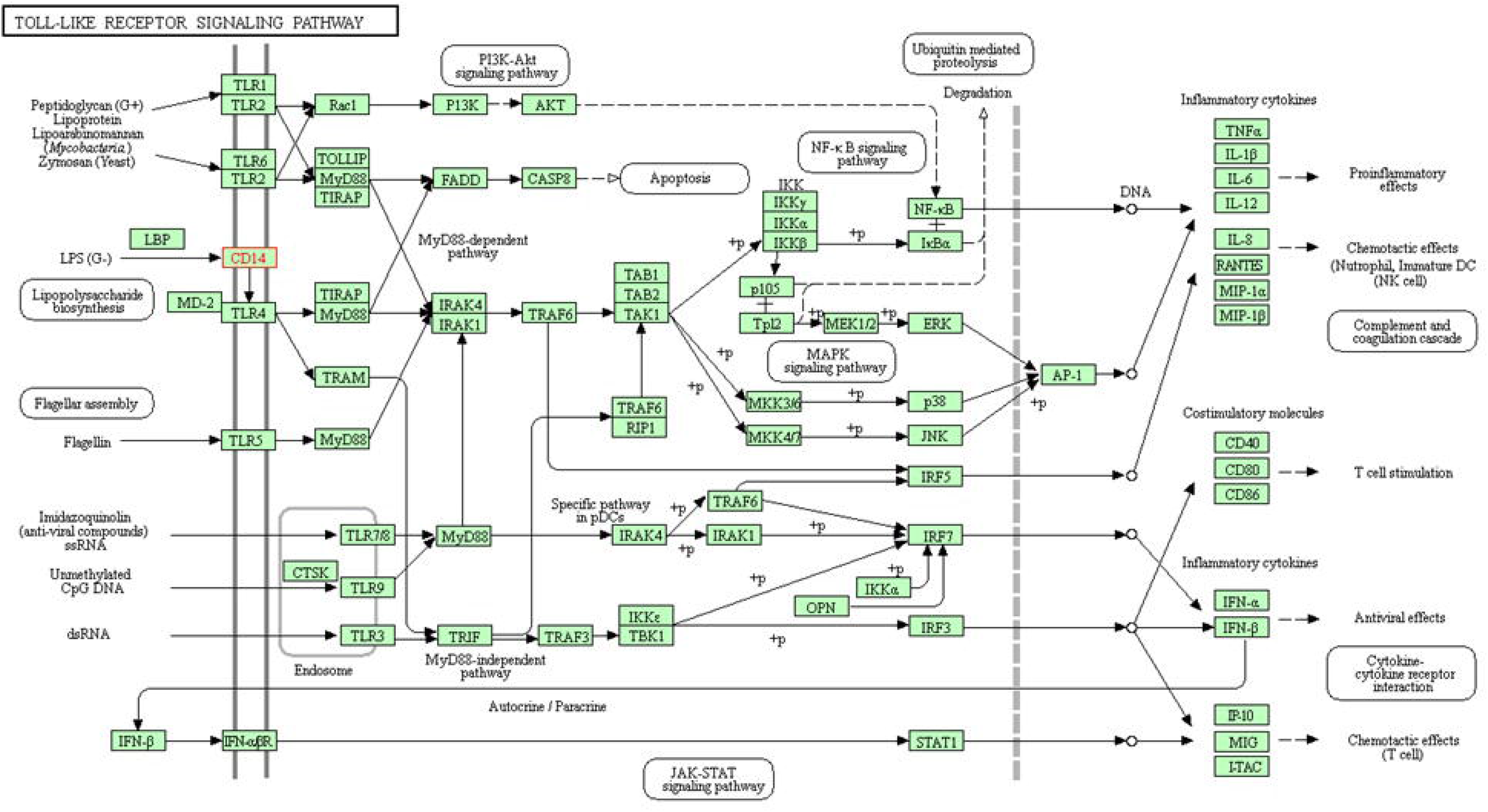
KEGG analysis depicting biochemical pathway involved in CD14 molecules.

### Molecular Phylogeny

Phylogenetic analysis of CD14 DNA sequences strongly suggests that avian diverge first from mammals with 4870 nucleotide substitutions with respect to CD14 gene (Fig 8). Rodents were the next to branch off (Fig 8). Phylogenetic tree for ruminants with other mammals and avian of both domestic and wild origin based on CD14 gene have been depicted in Fig 8. Ruminants were clustered together with about 300 nucleotide substitutions. Small ruminants as sheep and goat were clustered separately from large ruminants as cattle and buffalo. The closest ancestor for ruminants were found to be dog. Primates were genetically distant from ruminants at about 2400 nucleotide substitutions. Based on CD14 gene analysis, avian were observed to be genetically distant to mammals at about 4870 nucleotide substitutions. The genetic similarity/dissimilarity in terms of amino acid variations have been predicted in Table 3.

**Table 3:** Amino acid variations in different species of ruminants.

| Sl no. | Amino acid position | Buffalo | Cattle | Sheep | Goat | Remarks/ Domain present | Analysis by I-mutant | Analysis by Provean |
| --- | --- | --- | --- | --- | --- | --- | --- | --- |
| 1 | 13 | P | P | P | S | Site for signal peptide | Large decrease of stability |  |
| 2 | 14 | P | S | P | A |  | Large decrease of stability |  |
| 3 | 22 | T | T | T | K |  | Large decrease of stability |  |
| 4 | 23 | T | T | T | R |  |  |  |
| 5 | 30 | D | D | D | P | Domains for LPS binding site 1, LPS signaling site, site for LRR |  | Deleterious |
| 6 | 31 | D | D | D | Q |  |  |  |
| 7 | 32 | D | D | D | H |  |  |  |
| 8 | 74 | A | A | V | A |  | Large decrease of stability in A |  |
| 9 | 77 | N | T | D | D | LPS binding site | Large decrease of stability N, |  |
| 10 | 131 | M | T | T | T | LRR site |  |  |
| 11 | 143 | F | L | L | L | LRR site | Large decrease of stability |  |
| 12 | 173 | V | V | A | A |  |  |  |
| 13 | 179 | A | V | A | A |  |  |  |
| 14 | 186 | C | S | C | C | Site for di-sulphide bond | Large decrease of stability | Deleterious |
| 15 | 193 | E | G | E | E |  | Large decrease of stability |  |
| 16 | 195 | L | P | L | L |  | Large decrease of stability | Deleterious |
| 17 | 201 | S | F | S | S |  |  | Deleterious |
| 18 | 210 | G | R | G | G |  |  | Deleterious |
| 19 | 212 | M | R | M | M |  | Large decrease of stability |  |
| 20 | 221 | P | P | P | R |  | Large decrease of stability | Deleterious |
| 21 | 223 | L | L | L | P |  | Large decrease of stability | Deleterious |
| 22 | 236 | L | P | P | A |  | Large decrease of stability |  |
| 23 | 238 | G | G | G | R |  |  |  |
| 24 | 239 | V | V | V | P |  | Large decrease of stability | Deleterious |
| 25 | 248 | V | E | V | V |  | decrease of stability |  |
| 26 | 252 | S | S | S | N |  |  |  |
| 27 | 263 | A | L | A | A | Site for LRR | Large decrease of stability |  |
| 28 | 267 | A | C | A | A |  | Large decrease of stability |  |
| 29 | 276 | L | L | P | I |  | Large decrease of stability |  |
| 30 | 281 | L | L | L | C | Domain linker, leucine zipper | Large decrease of stability | Deleterious |
| 31 | 318 | E | E | E | Y | Domain Linker |  |  |
| 32 | 321 | D | D | V | V |  | Large decrease of stability |  |
| 33 | 336 | Q | Q | K | Q |  | Large decrease of stability |  |
| 34 | 337 | R | H | H | H |  |  |  |
| 35 | 345 | G | G | G | R | LRR |  | Deleterious |
| 36 | 349 | A | A | A | D | LRR | Large decrease of stability |  |
| 37 | 350 | C | C | C | R |  |  |  |
| 38 | 353 | S | F | S | P |  |  | Deleterious |
| 39 | 356 | T | T | T | V |  |  |  |
| 40 | 364 | A | A | A | V |  |  |  |
| 41 | 366 | L | L | L | Y |  | Large decrease of stability |  |

**Fig 8:**
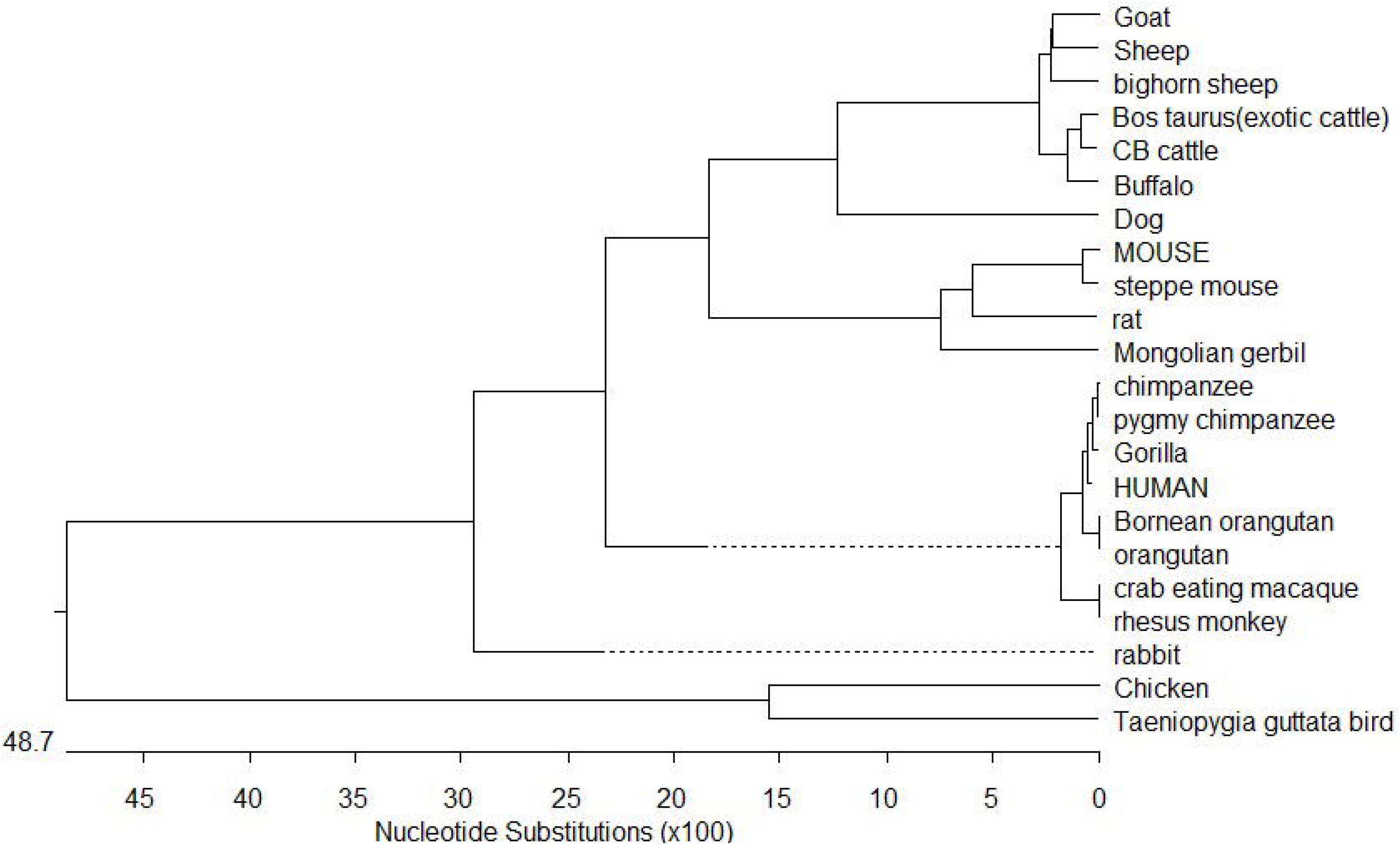
Phylogenetic analysis of mammals and birds with respect to CD14 gene

Trans-species polymorphism has been detected in ruminants (Table 4), where specific alleles were detected for small ruminants (goat and sheep) and large ruminants (cattle and buffalo). Twelve sites were detected (Table 4) with 5 synonymous and 7 non-synonymous nucleotide substitutions. Non-synonymous substitution exceeding synonymous indicates positive selection. Comparative sequence analysis of cattle and buffalo CD14 has been depicted in Table 4. Twelve non-synonymous nucleotide substitutions have been depicted in cattle and buffalo CD14 gene.

**Table 4:** Trans species polymorphism detected from molecular evolutionary study of CD14 gene for the ruminants.

| Nucleotide positions | Nucleotide in small ruminants (Sheep & Goat) | Nucleotide in large ruminants (Cattle and Buffalo) | Resulting amino acid change in small ruminants | Resulting amino acid change in large ruminants |
| --- | --- | --- | --- | --- |
| 102 | C | T | Synm |  |
| 181 | C | A | R | S |
| 194 | A | G | H | R |
| 204 | C | A | D | E |
| 223 | A | G | N | D |
| 229 | G | A | D | N (Buffalo)<br>T (Cattle) |
| 384 | T | C | Synm |  |
| 518 | C | T | A | V |
| 639 | G | A | Synm |  |
| 873 | A | G | Synm |  |
| 882 | G | C | Synm |  |
| 962 | T | A | V | D |

### Assessment of parasitic infestation in sheep

Screening of the faecal egg count (FEC) of the sheep (n=60) lead to study the health condition of the sheep which further prevailed us to divide them into two categories as Healthy and Diseased group. The mean FEC of the animal are given in the Table 5.

**Table 5:** Mean FEC of the Healthy and Diseased sheep.

| Health Status of the sheep | Mean Faecal Egg Count |
| --- | --- |
| Healthy | 50 ± 5.56 |
| Disease | 550 ± 9.45 |

### Differential mRNA expression profile for CD14 gene with respect to healthy and diseased sheep

The current study depicts increased expression profile of CD14 in diseased (H.contortus) infected sheep as depicted in Fig 9.

**Fig 9:**
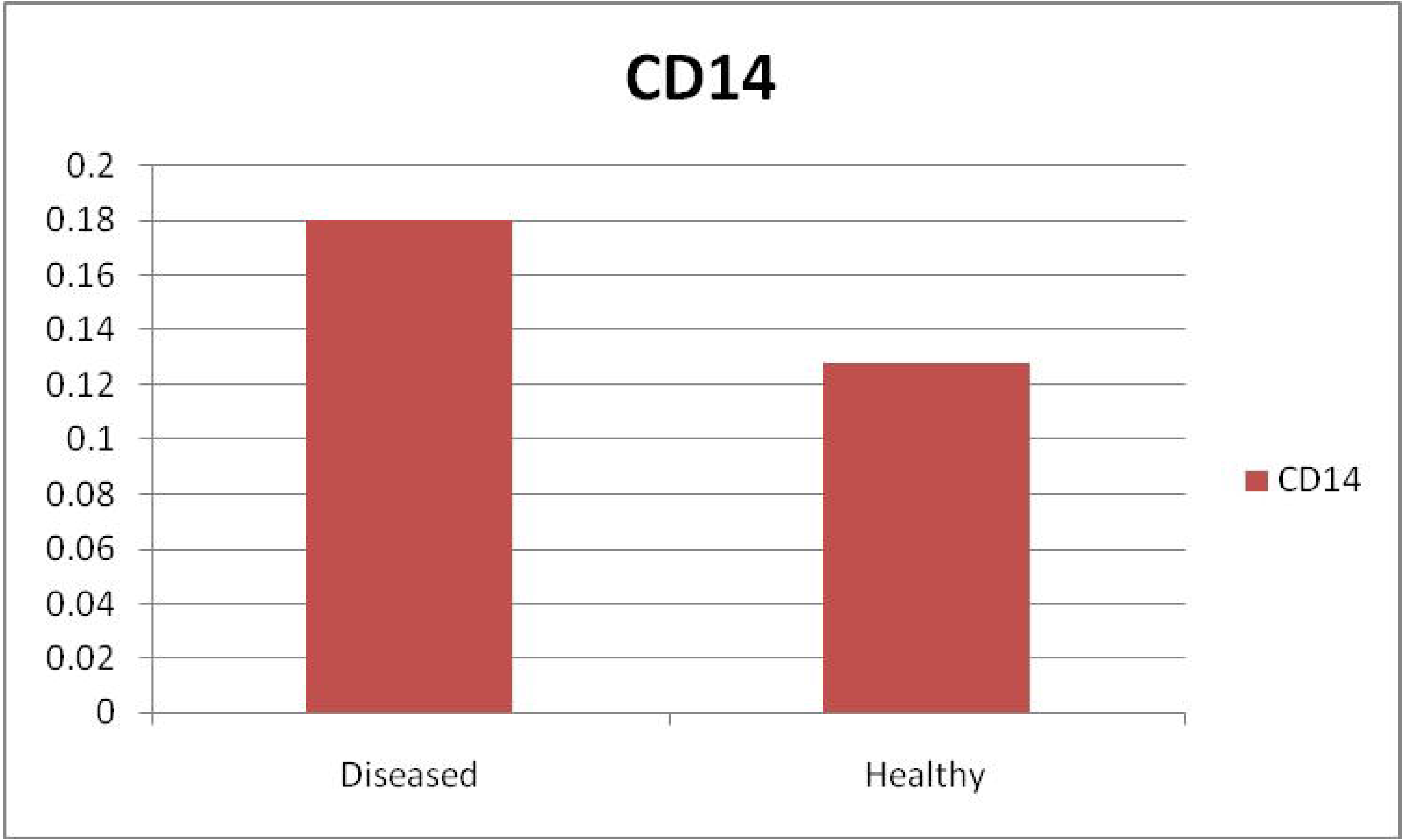
Differential mRNA expression profiling of ovine CD14 with respect to healthy and diseased (Haemonchus infected)

#### *In silico* study for the detection of binding site of CD14 with *Haemonchus contortus*

Molecular docking has revealed binding of CD14 with two identified surface molecule of *Haemonchus contortus* as alpha tubulin (Fig 10 A-B) and beta tubulin(Fig 10 C-D) with definite binding sites. Fig 10A depicts the structural alignment of alpha tubulin of H. contortus with CD14 gene, and aligned site being depicted as Glu27 (red sphere) and Ala284 (green sphere). Fig 10 B depicts the line diagram of the alignment of alpha tubulin of H. contortus with ovine CD14. Similarly Fig 10C depicts the structural alignment of beta tubulin of H. contortus with CD14 gene, and aligned site being depicted as Glu27 (hot pink sphere) and Ala284 (deep teal sphere). Fig 10 D depicts the line diagram of the alignment of alpha tubulin of H. contortus with ovine CD14.The study reveals distinct binding sites for CD14 with two most important structural proteins of Haemonchus contortus as alpha tubulin and beta tubulin. This finding indicates distinct role of CD14 in parasitic immunity. This is the first and novel report of the role of CD14 in parasitic immunity, hence comparison was not possible.

**Fig 10:**
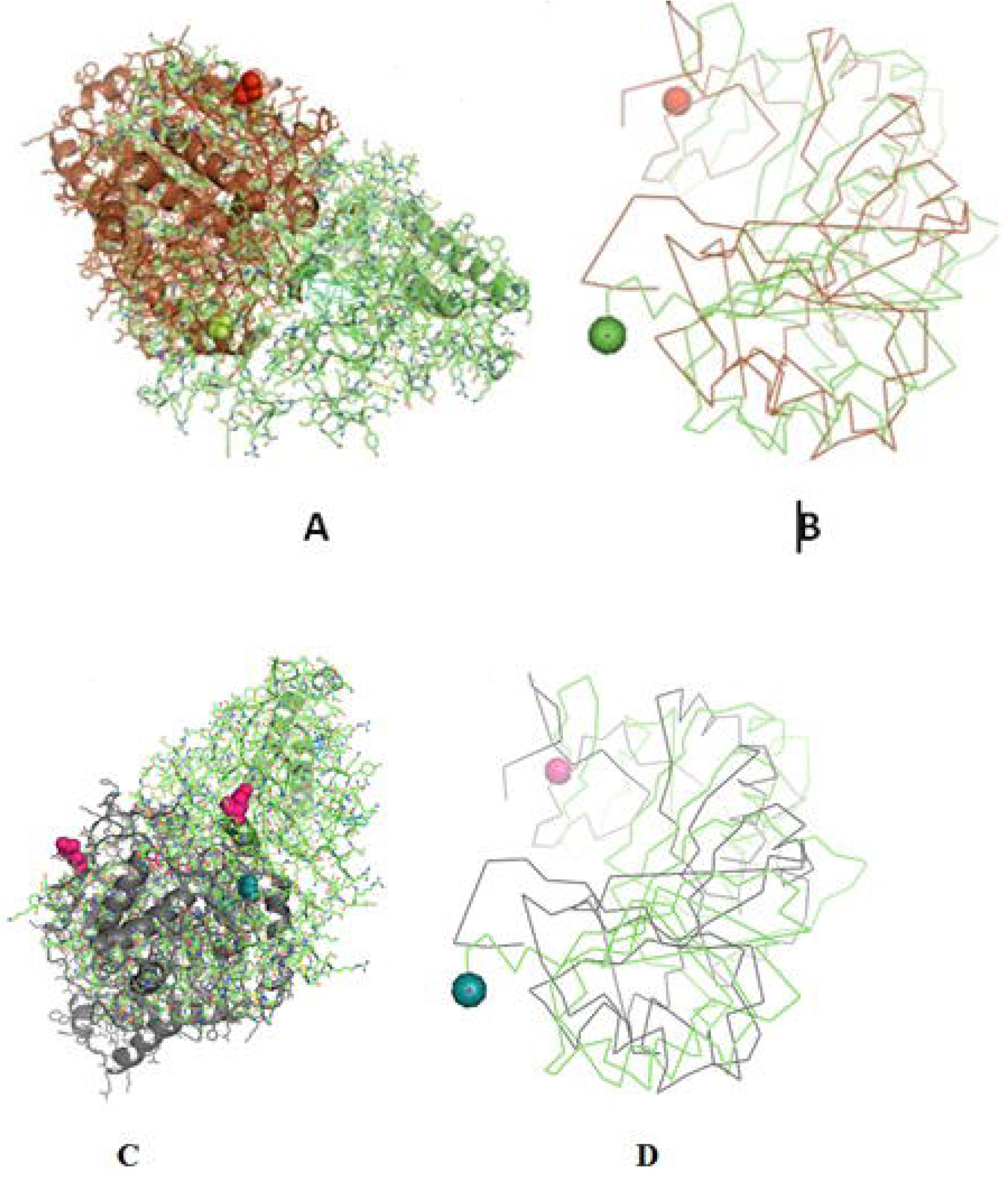
Binding of ovine CD14 with alpha tubulin and beta tubulin of *Haemonchus contortus*. A: Alignment of 3D structure of ovine CD14 molecule (green) with alpha tubulin (brown) of Haemonchus contortus B: Line drawing of binding site of CD14 molecule with alpha tubulin (brown) of Haemonchus contortus C: Alignment of 3D structure of ovine CD14 molecule (green) with beta tubulin (purple) of Haemonchus contortus D: Line drawing of binding site of CD14 molecule (green) with beta tubulin (purple) of Haemonchus contortus.

### Haematological and Biochemical parameters with respect to healthy and infected sheep

The hematological parameters were assesed with respect to diseased (H.contortus infected) vs. healthy sheep (Table 6). Significant differences were observed in Hb gm/dl, PCV%, TEC /cumm, TLC/cumm, Neutrophil %, Eosonophil%, Lymphocyte% . Better haemoglobin, PCV and TEC was observed in healthy sheep in comparison to that of infected. However, total leucocyte count was pronounced in infected sheep. A marked significant increase in eosinophil count was observed in infected sheep. Similarly pronounced increase was observed in neutrophil and lymphocytes, indicative of involvement of immune responsiveness.

**Table 6.** Haematological parameters in disease and healthy sheep.

| S.no | Parameter | Disease | Healthy |
| --- | --- | --- | --- |
| 1. | Hb gm/dl | 7.96<sup>b</sup>±0.19 | 10.37<sup>a</sup>±0.19 |
| 2. | ESR mm | 1.00±0.00 | 0.90±0.04 |
| 3. | PCV% | 27.84<sup>b</sup>±0.67 | 36.25<sup>a</sup>±0.64 |
| 4. | TEC /cumm (10<sup>6</sup>) | 3.98<sup>b</sup>±0.05 | 4.95<sup>a</sup>±0.08 |
| 5. | TLC/cumm | 9200.00<sup>a</sup>±159.23 | 6587.50<sup>b</sup>±210.81 |
| 5. | Neutrophil % | 53.50<sup>a</sup>±1.00 | 30.25<sup>b</sup>±0.59 |
| 6. | Eosonophil% | 16. 62<sup>a</sup>±0.49 | 2.12<sup>b</sup> ±0.44 |
| 7. | Basophil% | 0.00 | 0.00 |
| 8. | Lymphocyte% | 29.12<sup>b</sup>±0.29 | 60.87<sup>a</sup>±0.29 |
| 9. | Monocyte% | 1.37±0.18 | 1.87±0.29 |

The Mean values that have superscript, was showing significant difference between groups. The superscript 'a' was showing high mean value than 'b'.Significant at ( $p \leq 0.05$ ).

Significant differences were observed for healthy and infected sheep in terms of liver function test and kindney function test. Glucose level was observed to be better in healthy, while Creatinine, uric acid, urea, BUN level was better in infected sheep. Total protein, albumin, globulin and albumin: globulin ratio was observed to be better in healthy sheep (Table 7,8). While other reports indicate more in infected sheep (S.G.P.T. S.G.O.T, Alkaline Phosphatase, Total Bilirubin, Direct Bilirubin, Indirect Bilirubin(Table 7&8).

**Table 7.** Biochemical parameters (liver function test)in disease and healthy sheep.

| Sr. no | Parameters | Disease | Healthy |
| --- | --- | --- | --- |
| 1 | Total protein (g/dl) | 6.11<sup>b</sup>±0.02 | 7.77<sup>a</sup>±0.03 |
| 2 | Albumin (g/dl) | 1.92<sup>b</sup>±0.04 | 2.80<sup>a</sup>±0.03 |
| 3 | Globulin(g/dl) | 4.18<sup>a</sup>±0.06 | 4.97<sup>b</sup>±0.06 |
| 4 | Albumin:Globulin | 0.46<sup>b</sup>±0.01 | 0.58<sup>a</sup>±0.01 |
| 5. | S.G.P.T IU/L | 39.75<sup>a</sup>±1.4 | 32.12<sup>b</sup>±1.0 |
| 6 | S.G.O.T IU/L | 86.87<sup>a</sup>±2.6 | 68.87<sup>b</sup>±2.7 |
| 7 | Alkaline Phosphatase IU/L | 85.25<sup>a</sup>±3.89 | 64.87<sup>b</sup>±2.09 |
| 8 | Total Bilirubin mg/dl | 0.67<sup>a</sup>±0.07 | 0.37<sup>b</sup>±0.03 |
| 9 | Direct Bilirubin mg/dl | 0.30<sup>a</sup>±0.02 | 0.16<sup>b</sup>±0.01 |
| 10 | Indirect Bilirubin mg/dl | 0.37<sup>a</sup>±0.09014 | 0.21<sup>b</sup>±0.05 |

The Mean values that have superscript, was showing significant difference between groups. The superscript 'a' was showing high mean value than 'b'.Significant at ( $p \leq 0.05$ ).

**Table 8.** biochemical parameters (kidney function test) in disease and healthy sheep.

| S.No. | Parameters | Healthy | Diseased |
| --- | --- | --- | --- |
| 1 | Glucose mg/dl | 55.50<sup>b</sup>±1.21 | 68.62<sup>a</sup>±2.33 |
| 2 | Creatinine mg/dl | 1.71<sup>a</sup>±0.05 | 1.20<sup>b</sup>±0.02 |
| 3 | Uric acid mg/dl | 0.89<sup>a</sup>±0.01 | 0.76<sup>b</sup>±0.04311 |
| 4 | Urea mg/dl | 48.37<sup>a</sup>±1.2 | 36.12<sup>b</sup>±1.56 |
| 5 | BUN mg/dl | 23.62<sup>a</sup>±0.73 | 16.87<sup>b</sup>±0.95 |
| 1 | Glucose mg/dl | 55.50<sup>b</sup>±1.21 | 68.62<sup>a</sup>±2.33 |
| 2 | Creatinine mg/dl | 1.71<sup>a</sup>±0.05 | 1.20<sup>b</sup>±0.02 |

## Discussion

Ovine CD14 was characterized and the important domains were identified in comparison to other ruminant species. GC content of CD14 gene of human (62.94%), dog (64.07%) was found to be more than ruminants while rodent had less GC content, e.g. for rat (56.12%). High GC content was observed to be an inherent characteristic of CD14 gene. Direct relationship of the length of the coding sequence was observed with GC content of the gene (Pozzoli 2008). Species-wise differentiation in GC content may explain the variability in the stability of CD14 gene among the species. It is known that nucleotide Guanine (G) and cytosine (C) are bounded by triple hydrogen bond in double helix DNA. Thus, it is evident that GC bond is stronger than AT bond. Similarly GC content is reported to have significant effect on genome functioning and species ecology (Smarda et al., 2014). The importance of GC rich isochores has been predicted. The thermostability, bendability, ability to B-Z transition and curvature of DNA helix was observed to be correlated positively with GC content (Vinogradov, 2003).

Alignment of the derived peptide sequence of ruminants revealed that leucine residues were mostly conserved across the species (Kim et al., 2005), except at amino acid position 143. Cysteine residue of CD14 was found to be conserved across ruminant species and other species (Fig 4). The CD14 molecule found in ruminants was predicted to contain the higher percentage of leucine (Table 2), which is similar to mouse (17.66%) and human (15.5%) (Setuguchi et al., 1989).

Due to protein folding, certain grooves are developed in CD14, which in turn are responsible for the development of a binding site for ligands. Since the models are developed through homology modeling, 5th disulphide is not visualized in the model. Similar five disulphide bonds were also observed in mice (Kim et al., 2005) and human (Meng et al., 2008). Cys287 is tyrosine moiety in the mouse CD14 sequence, hence from the homology modeling, the fifth disulphide could not be detected.

The crystal structure of CD14 was observed to contain 5 disulphide bonds both in human (Meng et al., 2008) and Mus musculus CD14 molecule (Kim et al., 2005). Biochemical studies have also detected five disulfide bonds in human (Setuguchi et al., 1989). The impact for disulfide bonds differ in human CD14 folding. Out of five disulfide bonds, first and second disulfide bonds were observed to be indispensible. The third and fourth disulphide bonds were reported to be important, while the fifth or the last (Cys287–Cys333) was dispensable (Meng et al., 2008). Disulfide bond have greater effect on CD14 structure and integrity, as in case of mouse. The first and second disulfide bond were observed in the β sheets in the inner concave surface (the “core” structure), whereas the third and fourth disulfide bonds were observed in the loops and helices on the outer surface (the “peripheral” structure).The first disulphide bond is responsible for the structural and functional integrity of CD14 molecule. It was inferred that CD14 folding was not determined by third, fourth and last disulphide bonds. The substitutional mutation had been observed in the 3rd and 4th disulphide bond. Cysteine was substituted with alanine, but the function of CD14 is not altered (Meng et al., 2008).

In ruminants, four LPS binding site and three LPS signaling sites were predicted. The generous size of the pocket of the CD14 receptor may allow structural variation in the hydrophobic portion of the ligand, however, the hydrophobic bottom and walls of the pocket are rigid. Hydrophilic part of the ligand has structural diversity. This is due to the considerable flexibility of the hydrophilic part of the rim leading to the development of multiple grooves for ligand binding. Similar LPS binding site and LPS signaling site were reported in mice (Kim et al., 2005) and human (Setuguchi et al., 1989). LPS signaling site also overlaps with LRR. It has also been observed that the first LRR (Leucine-rich repeat) overlaps with LPS-binding site region-I. CD14 is a pattern recognition receptor and are rich in leucine-rich repeat and CD14 are rich in leucine moiety as evident by high leucine percentage. Ligand specificity or flexibility of the rim of CD14 is important for pathogen binding site for a wide range of pathogens. There was the report of three, five, four and five glycosylation sites for N-linked glycosylation in Bos taurus (Fillip et al., 2001), Mus musculus (Setuguchi et al., 1989), Homo sapiens (Setuguchi et al., 1989), Rattus norvegicus (Takai et al., 1997) respectively. N-linked glycosylation is vital for the molecule as it supports the molecule to be present either in membranous or soluble form. O-linked glycosylation site for ruminant CD14 differs, three in Buffalo and five in both goat and crossbred cattle (Table 2), but absent in human (Savedra et al., 1996). O-linked Glycosylation is needed when hydrophilic clusters of carbohydrates alter the polarity and solubility of protein or protein folding. *Homo sapiens, Mus musculus,* were reported to have 10 LRR each. A trend was observed for the relationship of number of LRR and disease susceptibility. Pathogen recognition is an important phenomenon in any innate immune response. LRR for CD14 in extracellular domain has a major role in pathogen recognition.

Similar findings as ovine CD14 for GPI anchor were reported in other species like human and mouse. GPI anchor is an important phenomenon for any membranous protein. GPI anchor for CD14 is important for bridging between CD14 and cell surface. In avian, CD14 was observed to be transmembranous, not GPI anchored (Wu et al., 2009). Amino acid positions responsible for NES are 12, 15, 16, 17, 117, 122 127 in ruminant CD14. Since nuclear export signal (NES) aids in export of the CD14 peptide from nucleus to the cytoplasm via the nuclear pore complex (Cour et al., 2004), there is possibility of impairment of this function in goat CD14. In terms of alpha helix and Beta strandsof ovine CD14 has similarity in mice containing seven number of alpha helix, while differs in Beta helix . Mice CD14 contains 13 Beta helix in contrast to eleven in ruminants. It seems that Beta helix have a relationship with the innate immunity of CD14 molecule. Nine Beta strand starting from β2, β4 to β11 has an overlapping with leucine rich repeats (LRRs). LRR in turn have role on LPS binding and LPS recognition site, as already discussed, which forms one of the basis for innate immunity.

Since CD14 molecule forms its functional form by dimerization, leucine zipper leads to dimerization. Leucine zippers, a common three-dimensional structural motif, (64) were observed to be responsible for gene expression, since these were an integral component of DNA-binding domain in a variety of transcription factors. The presence of leucine zipper generates adhesion forces in parallel alpha helices (Landschulz et al., 1988) and this region was found to be conserved in ruminant CD14.

Domain linker sequences connect two structural domains and in turn act as a scaffold and prevent unfavourable interactions between folding domains (George and Heringa, 2002).

CD14 molecule was predicted to be mostly expressed on cell membrane or cell surface for the ruminants under study, which justifies it to be a receptor molecule. Two forms of CD14 as membranous and soluble form are in existence (Paape et al., 1996, Wright et al., 1991). Cloned ovine CD14 cDNA in current study was predicted to be existing in membranous form since it contained GPI anchor (Paape et al., 1996, Wright et al., 1991).

The finding is similar to study conducted by (Kim et al., 2005), who studied crystal structure of Mus musculus CD14 molecule.

Comparison of ovine CD14 with that of goat has revealed certain differnces. Epidemiological studies also depict that sheep were better resistant to goat in terms of infectious and parasitic diseases. GC content were less in sheep. It has been reported that there is preferential fixation of GC alleles leading to biasness of GC nucleotides over AT during DNA repair. GC content is important is respect of evolving recombination rate, regulated replication or expression timing, bending ability of DNA and to B-Z transition, transcriptional efficiency as already explained earlier.

It is evident that higher mutation frequency was detected in goat (0.56), with mutations detected at 25 amino acid positions, out of which thirteen sites were observed to be thermodynamically unstable. Seven sites were observed to be deleterious in nature.

Mutations detected as P13S, P14A in goat are involved in domain for signal peptide was observed to have less structural stability. Mutations were observed as D30P, D31Q, D32H, which are responsible for important domains as LPS binding site1, LPS signalling site and LRR site, where D30P was observed to be deleterious. In site 1 of LPS binding site, there is drastic shift of acidic amino acid to basic, as Aspartic acid to proline, aspartic acid to glutamine, Aspartic acid to histidine. Since there is a change of polarity, the pathogen binding activity is also adversely effected.

P276I was observed to be thermodynamically unstable, but this site constitutes LRR, which is important for LPS signalling. Thus this mutation may cause depressed function of pathogen binding activity of CD14 of goat in contrast to sheep. Another important site for deleterious mutation is L281C, which are responsible for the domain linker, leucine zipper. K336Q was thermodynamically less stable in goat CD14 and is responsible for the site of domain linker.

A cluster of mutations were observed in goat CD14, which are important for LRR site, ultimately effecting pathogen binding and signalling, which are G345R, A349D, C350R, S353P,T356V, A364V, L366Y.

MD2 is an important component of TLR4 signaling complex. Members of the MD-2–related lipid-recognition protein super- family contains two beta sheets arranged in ab cup fold generating a centralized hydrophobic LPS-binding cavity (Inohara et al., 2002). CD14 sensitizes cells to LPS through the process of delivering this lipid moeity to MD-2. The hydrophobic pocket is occupied mostly by four acyl chains of the ligand buried inside for the complex of both mouse and human CD14 with MD-2 bound to lipid IVa (Ohto et al., 2007, Ohto et al., 2012). TLR4/MD-2/LPS homodimeric signaling complex depicts the hydrophobic binding pocket sequestering five of the six fatty acid chains. The rest of the unbound acyl chain lies along the surface of MD-2 and, along with the F126 loop of MD-2, generates a new hydrophobic patch promoting homodimer formation through the association of TLR4/MD-2/LPS complex (Teghanem et al., 2007).

LPS-binding pocket (hydrophobic) was predicted at the amino terminal end through a homology based study with 3D structure of CD14 of Mus musculus (Kim et al., 2005) and Homo sapiens (Setuguchi et al., 1989). CD14 molecule, α-helices were located at the convex surface, and β strand were located at the concave surface of horse-shoe shaped CD14 structure, represented in Figure 3 & 5. N terminal hydrophobic pocket revealed a conservation of pocket size and hydrophobicity. Asparagine ladder with conserved asparagines residue had been observed, which is a structural feature for leucine rich repeats. The variable residues in LRR are hydrophilic and exposed to concave surface of horseshoe structure. Some of the similar studies had been conducted earlier in ruminants (Pal and Chaterjee, 2009, Pal et al., 2011, Pal et al., 2013, Pal et al., 2014).

CD14 has been considered as an important gene to explore a wide range of open questions in evolutionary ecology, since a wide range of nucleotide variability have been obse0rved with mutational hot spot detected. It was attempted to identify the Trans species polymorphism (TSP).

TSP generally occurs by passage of alleles from ancestral to descendant species. Neutral alleles may be shared by species which are genetically similar. Balancing selection has been regarded as the major evolutionary force responsible for long lasting TSP. Similar TSP as identified in ruminants with respect to CD14 gene have also been identified in MHC Class II gene (Klein et al., 1998, Li et al., 2011). Speciation is a complex process which involves TSP, divergence of allelic lineages at the coding sites for both synonymous and non-synonymous mutations. These processes involve millions of years ago. (Klein et al., 1998).

Pronounced expression of CD14 was observed in diseased sheep compared to that of healthy sheep. As per our literature search, it is the first report in sheep. Similar better expression profile of CD14 was also detected in mice infected with another parasite as *Schistosoma mansoni* (Tundup *et al*., 2014).

Some more studies suggested that CD14^+^, CD16^+^ cells help to control parasitic burden and play important role in resistance to *P. vivax* infection in human (Antonelli *et al*., 2014). However, in our present study increased expression of CD14 gene in diseased sheep may be due to its action as pattern recognition receptor in parasitic infection. It was reported that parasite-derived E/S product was detected by CD14^bright/^CD16^+^ intermediate monocytes in case of schistosome-infected patients compared to that of uninfected individuals (Turner et al., 2014). In addition, CD14 has a significant role in regulating the immunity in *S. mansoni*-infected mice, through the induction of both TH1 and TH2 through macrophage M1/M2 phenotype. CD14^bright^CD16^+^ intermediate monocytes are major producer of IL-10 (Skrzeczynska-Moncznik J *et al*.,2008). The Th1 and Th2 cell released different kinds of cytokines such as interleukin (IL-2), interferon-gamma (IFN-γ) and tumour necrosis factor-alpha (TNF-α) from TH1 cells and IL-10 produce from Th2 type cells and these cytokines are responsible to stimulates B cell differentiation, production of different immunoglobulin such as IgE, IgG1, IgG4 and IgA, mastocytosis, and eosinophil activation and function ( Janeway *et al.,* 2004).

IL-10 leading to eosinophil mobilization, intestinal mast cell accumulation and production of IgE, mucosal mast cell infiltration, intestinal eosinophilia, elevated serum IgE, increased level of parasite specific IgG-1 which are responsible to eradicate worms (Tizzard,2004).

Similar studies that correspond to our finding shows marked increase in bood eosinophil and abomasal tissue, degranulation of eosinophil and it releases some peroxidases and basic protein and cationic protein which are cytotoxic to helminth.(F.Alba-Hurtado *et al*.,2013).

## Conclusion

CD 14, an important gene for innate immunity has revealed immense variability between species, which may lead to variability in resistance to diseases. This variability in disease resistance may be due to variability in ligand binding ability of the CD14 molecule. The greater susceptibility of goat compared to sheep, and greater susceptibility of cattle compared to buffalo revealed mutations in LPS binding site of goat/ cattle CD14 molecule, which causes altered polarity, decreased thermodynamic stability and the deleterious mutation. Since hypervariable site or mutational hot spot has been detected in CD14 gene, due to greater genetic variability, evolutionary study is very promising. Phylogenetic analysis reveals that ruminants were evolutionary closer, but transpecies polymorphism has been detected among the ruminants. The distinct role of CD14 in antiparasitic immunity is reported here for the first time through *in silico* molecular docking and later confirmed through differential mRNA expression profile with QPCR for healthy and infected samples.

## Acknowledgement

The authors like to thank Department of Biotechnology, Ministry of Science and Technology, Govt. of India for providing the financial support for carrying out the research work (Grant No. BT/Bio-CARe/04/10100/2013-14). The funders had no role in study design, data collection and analysis, decision to publish, or preparation of the manuscript. The authors are equally thankful to West Bengal University of Animal and Fishery Sciences and Indian Veterinary Research Institute for carrying out the work.

## Conflict of Interest

It is to declare that there exists no conflict of interest.

## Notes

### Competing Interest Statement

The authors have declared no competing interest.

